# Ribozyme-phenotype coupling in peptide-based coacervate protocells

**DOI:** 10.1101/2022.10.25.513667

**Authors:** Kristian Le Vay, Elia Salibi, Basusree Ghosh, T-Y Dora Tang, Hannes Mutschler

## Abstract

Condensed coacervate phases are now understood to be important features of modern cell biology, as well as valuable protocellular models in origin of life studies and synthetic biology. In each of these fields, the development of model systems with varied and tuneable material properties is of great importance for replicating properties of life. Here, we develop a ligase ribozyme system capable of concatenating short RNA fragments into extremely long chains. Our results show that formation of coacervate microdroplets with the ligase ribozyme and poly(L-lysine) enhances ribozyme rate and yield, which in turn increases the length of the anionic polymer component of the system and imparts specific physical properties to the droplets. Droplets containing active ribozyme sequences resist growth, do not wet or spread on unpassivated surfaces, and exhibit reduced transfer of RNA between droplets when compared to controls containing inactive sequences. These altered behaviours, which stem from RNA sequence and catalytic activity, constitute a specific phenotype and potential fitness advantage, opening the door to selection and evolution experiments based on a genotype – phenotype linkage.

## Introduction

Many biological biomolecular condensates are formed from RNA and proteins or peptides^1,2^, and coacervate phases formed by charge-mediated phase separation are part of the mechanism that drives the development of membraneless organelles in modern biology^3,4^. Condensed phases studied in the context of origin of life can also be formed from RNA and peptides. Catalytic nucleic acids, including ribozymes and DNAzymes, are central to ‘Nucleic Acid World’ hypotheses, where they act as both the medium of information storage and the catalyst for its replication in early life-like systems. Peptide-RNA condensation forms discrete protocellular compartments^5,6^. RNA-peptide interactions are in many cases beneficial for the function of these early catalysts^7,8^, and may promote the folding and oligomerisation of certain peptides^9^. Perhaps surprisingly, the function of ribozymes and other nucleic acid enzymes in coacervate phases has only been recently established, with catalytic rate being enhanced and enzyme function altered in some cases^10–14^. For example, coacervation with poly(L-lysine) shifts the reaction equilibrium of a minimal hairpin ribozyme from cleavage to ligation^15^.

The ability of coacervate phases to strongly partition a wide range of molecular and macromolecular species^16^, and to support a variety of complex enzymatic processes^17^, makes them appealing proto- and artificial cell models in origin of life studies^18^, synthetic biology^19^ and modern biology^20^. Simple coacervates that form from oppositely charged polymers have been rigorously investigated for their ability to selectively partition biomolecules and host a variety of different chemistries and reactions *in vitro*^20,21^. However, biological condensates formed by protein-protein or protein-nucleic acid interactions are dynamic systems, and their formation, dissolution and physical properties are subject to spatiotemporal regulation^22^. Similarly, in order to realise a truly convincing model proto- or artificial cell, life-like behaviours such as growth, division and other dynamic, responsive or non-equilibrium processes are essential. Several dynamic and responsive coacervate systems have been established, which are characterized by a phase change in response to environmental stimuli such as light exposure^23–25^, changes in temperature^26^ or pH^27,28^. Furthermore, non-equilibrium environments formed by gas bubbles inside heated rock pores have been shown to drive the growth, fusion, and division of otherwise inert coacervate microdroplets^29^.

Enzymatic processes that alter the properties of the coacervate components also affect coacervate properties and behaviour. In charge-based condensates, inducing phase change via enzymatic processes has been achieved by, for example, the conversion of ADP into charge-dense ATP^30^, or the alteration of peptide charge state by phosphorylation^31^. This has allowed the reversible generation of coacervate droplets by enzymatic networks. In addition, the polymerisation of UDP in U_20_-spermine coacervates by polynucleotide phosphorylase has been shown to induce transient non-spherical coacervate morphologies^32^. All these systems depend on the action of proteinaceous enzymes, which presumably emerged later in molecular evolution, perhaps after the first protocells. However, droplets capable of dynamic change via the action of nucleic acid enzymes such as ribozymes have not been previously reported, partially because the catalytic repertoire of these catalysts is limited when compared to their proteinaceous counterparts^33^. Beyond environmental factors such as solution pH and salt concentration^34^, the physical properties and association behaviour of coacervate droplets are determined by factors such as component chain length^35^ and charge density^36^. Of these factors, we noted that polymer chain length could potentially be addressed by the action of a nucleolytic or ligase ribozyme. Thus, in a coacervate composed of RNA and a positively charged polymer, ribozyme catalysed RNA cleavage or ligation might alter the physical properties or association behaviour of the system if a sufficiently large change in average RNA length were achieved. In particular, we hypothesised that the elongation of the RNA component could lead to the formation of a denser separated phase with altered physical properties^34,37^.

In this work, we demonstrate that coacervate microdroplets formed from a ligase ribozyme and lysine-based peptides display reciprocal behaviour, in which coacervation enhances and modulates ribozyme activity, and ribozyme activity modulates droplet properties and therefore phenotype. We employ a modified ligase ribozyme that ligates short substrate strands into long concatemers, thus increasing the length of the polyanionic coacervate component. We find that coacervation enhances the rate of substrate ligation 50-fold and inhibits the formation of circular reaction products. In turn, the activity of the ribozyme imparts specific physical properties or phenotypes to the droplets it is contained within, which are not observed in droplets containing inactive ribozymes. These altered behaviours include the inhibition of droplet growth, surface wetting and content exchange, which could provide fitness advantages under certain conditions. Connecting the sequence information of the ribozyme RNA to the physical properties of the resulting microdroplets is an example of a phenotype-genotype linkage, which is a fundamental requirement for the Darwinian evolution of protocellular systems^38^.

## Results

To investigate the effect of ribozyme activity on coacervate behaviour, we initially sought to design a ribozyme system capable of increasing RNA chain length via concatenation. Although several examples of RNA ligase ribozymes have previously been reported^39–41^, these typically catalyse the ligation of a single junction, which results in only a modest overall increase in average RNA chain length. To achieve greater product lengths, we harnessed the catalytic core of the R3C ligase ribozyme (E_R_), whose RNA ligation activity is based on 5’-triphosphate activated substrates^39^. This system has recently been shown to be active in the presence of poly(L-lysine) under certain conditions, and so was a promising starting point when considering compatibility with coacervate systems^14^. We previously redesigned the ribozyme-substrate complex to iteratively produce long RNA concatemers from short oligonucleotides^42^. In our final design, the ribozyme (E_L_) catalyses concatenation of a 31 nt substrate (**Figure 1a**). Screening of reaction conditions established that strong activity was observed at pH 8.6 in the presence of 10 mM MgCl_2_ at a range of temperatures (30, 37 and 45 °C) **(Figure 1 – supplement 1)**. The strongest activity was observed with a 1:1 monomer concentration ratio of substrate:ribozyme, while excess substrate inhibited the formation of longer length products at lower temperatures, most likely because the ribozyme has a higher probability of binding unligated substrates than already growing chains. In order to reduce hydrolytic degradation of RNA, we chose to maximise activity at 30 °C with a 1:1 substrate:ribozyme ratio in all subsequent experiments.

**Figure 1.**
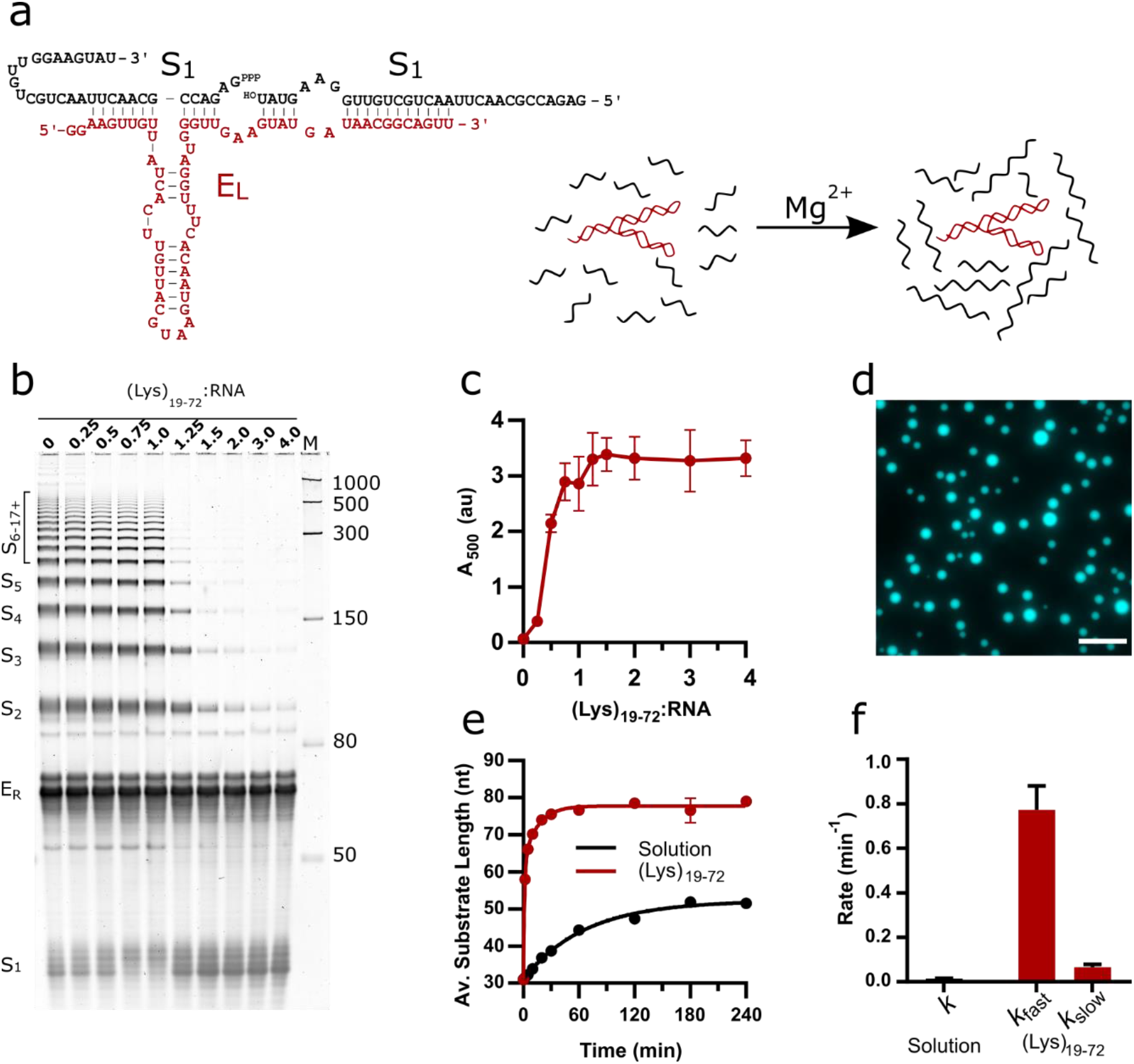
Design of the EL R3C ladder ribozyme system and activity in the presence of (Lys)_19-72_ peptides. **a)** The structure of the ladder ribozyme and a schematic showing its function. The ribozyme is shown in red, whilst the substrate strands are shown in black. **b)** An 8 % urea PAGE showing the products of the R3C ladder system in solution and varying ratios of (Lys)_19-72_ to R3C RNA (total monomer concentration = 1 mM) after a 2 h reaction at 30 °C, pH 8.6 and 10 mM MgCl2. **c)** Variation in absorbance at 500 nm upon addition of varying ratios of (Lys)_19-72_ to the EL RNA before ligation. Data points are an average of three technical replicates. Error bars are standard deviations. **d)** Example fluorescence microscopy image of (Lys)_19-72_:RNA condensates at a ratio of 0.75:1 (Lys)_19-72_:RNA, imaged using 10 % Cy5-tagged substrate strand. Scale bars = 20 μm. **e)** Kinetics of chain elongation in solution (black, first order model), and with 0.75:1 (Lys)_19-72_:RNA (red, second order model) at 30 °C, pH 8.6 and 10 mM MgCl_2_. Data points are an average of three technical replicates. Error bars are standard deviations. **f)** Chain extension rate constants for the R3C ladder ribozyme in solution (black, first order model) and in the presence of 0.75:1 (Lys)_19-72_:RNA (red, second order model). Equivalent data for condensates formed from the shorter (Lys)_5-24_ peptide is shown in **Figure 1 – supplement 2**.

The phase separation behaviour of the ribozyme system in the presence of poly(L-lysine) was initially investigated by titrating increasing amounts of (Lys)_19-72_ into a fixed concentration of RNA (total monomer concentration = 1 mM). The activity of the ribozyme was inhibited in the presence of excess peptide ((Lys)_19-72_:RNA > 1) (**Figure 1b**). The reported concentration ratios are calculated from RNA and peptide monomer unit concentrations, and as such are also equivalent to charge ratios. The addition of (Lys)_19-72_ led to a gradual increase in turbidity due to phase separation above the critical coacervation concentration of CCC_19-72_ ≈ 0.14:1 (Lys)_19-72_:RNA **(Figure 1c)**. These experiments were repeated with a shorter peptide ((Lys)_5-24_) (**Figure 1 – supplement 2**), for which the onset of coacervation occurred at higher peptide:RNA ratios (CCC_5-24_ ≈ 0.93:1 (Lys)_5-24_:RNA), corroborating previous observations with poly(L-lysine) and the HPz ribozyme^15^. In this case, the activity of the ribozyme was not inhibited in the presence of excess peptide. For further experimentation, we selected specific peptide:RNA ratios of 0.75:1 (Lys)_19-72_:RNA and 3:1 (Lys)_5-24_:RNA. Whilst these two conditions are not directly comparable due to their differing concentration ratios, the difference in critical coacervation concentration between the two peptide lengths meant that no single ratio supported both droplet formation and ribozyme activity in both systems. Both points occur shortly before the respective maxima in the peptide titration turbidity curve, allow the formation of liquid coacervate droplets and do not suppress ribozyme activity (**Figure 1c and Figure 1 - supplement 2b**). Fluorescence imaging of samples at these ratios confirmed the formation of phase separated coacervate droplets that strongly partitioned the Cy5-labelled RNA substrate (**Figure 1d and Figure 1 - supplement 2c**).

The kinetics of chain elongation and final product length at these ratios was greatly enhanced in the coacervate phase compared to solution (**Figure 1e-f**). The kinetic analyses show that the addition of either (Lys)_n_ peptide resulted in an approximately 50-fold increase in the rate of concatenation and led to the formation of substrate chains with an average length approximately 30 nt greater than those produced in solution (**Figure 1e)**. In solution, the kinetics of chain elongation were best approximated using a first order model (*k* = 1.5 × 10^−2^ ± 1.0 × 10^−3^ min^−1^), and a final average product length (n) of 52.2 ± 0.5 nt was observed (**Figure 1e**). In the (Lys)_19-72_ coacervate phase, the ribozyme kinetics were best described by a two-phase model (*k*_fast_ = 7.7 × 10^−1^ ± 6.6 × 10^−2^ min^−1^, *k*_slow_ = 6.6 × 10^−2^ ± 1.3 × 10^−2^ min^−1^), with a final product length of n = 77.8 ± 0.4 nt for (Lys)_19-72_. Similar values were obtained for the short peptide (**Figure 1 – supplement 2e, source data**). We have previously observed a coacervation-induced shift from monobasic to biphasic kinetic behaviour for the hammerhead ribozyme, albeit without an associated increase in rate^10^. The increase in rate observed here is most likely due to the high concentration of ribozyme and substrate that occurs in coacervates directly formed from catalytic RNA and peptides^15^.

In solution, the E_L_ ribozyme produced both linear and circular concatenates, the latter of which are visible on the urea – PAGE gel as additional bands visible in between and above the regularly spaced linear ladder of products, due to the different mobility of linear and circular products through the gel matrix. The presence of circular products was confirmed by treatment of reacted RNA with RNAse R, a 3’ to 5’ exoribonuclease that only digests linear strands **(Figure 1 – supplement 3)**. The presence of the long peptide (0.75:1 (Lys)_19-72_:RNA) suppressed circularization altogether, whilst the short peptide (3:1 (Lys)_5-24_:RNA) reduced the formation of circular products.

Having demonstrated that the E_L_ ribozyme is capable of substrate concatenation in both solution and in (Lys)_n_ coacervate droplets, and that its activity is enhanced within the coacervate environment, we aimed to determine if the elongation of the RNA component would lead to differences in droplet properties, thus altering phenotype. To identify changes due solely to chain concatenation, we developed an inactive mutant of the ribozyme by introducing several point mutations **(Method section - tables 1 and 2)**, without altering the substrate binding region. Ribozyme activity assays revealed that the mutant was completely inactive in solution and in the presence of both (Lys)_n_ peptides at the previously specified charge ratios **(Figure 2 – supplement 1)**. Populations of droplets containing either the active or inactive ribozyme were loaded into a passivated well plate and imaged over the course of 24 h. In these experiments, (Lys)_n_ was added to the RNA reaction mixture immediately, so that all ligation activity would occur in the presence of the peptide. In all cases, phase separated droplets were observed which persisted over the course of the experiment **(Figure 2a, c and Figure 2 - supplement 2a, a1c)**. Whilst many droplets remained static, some coarsening to form larger droplets was observed, particularly in the droplet populations containing the inactive ribozyme. To fully quantify these observations, a segmentation algorithm was used to identify droplets and thereby measure their size and population density^43^. Droplets which contained active ribozyme maintained a constant average area over the 24 h period, whilst droplets containing inactive ribozyme grew, in some cases reaching a final area more than twice that of the active system **(Figure 2e and Figure 2 – supplements 2 and 3)**. As the average droplet area increased for the inactive droplets, there was a concomitant decrease in droplet population density (the average number of droplets per 100 μm^2^), indicating that the growth was at least, in part, due coalescence events between the droplets **(Figure 2f)**. A minimal decrease in droplet population density was observed for active droplets, suggesting that lack of coalescence prevented growth. Similar behaviour was observed for both the long and short peptide **(Figure 2 – supplements 2 and 3)**, although for the short peptide system, the number of both active and inactive droplets decreased similarly over the course of the experiment, indicating coalescence was not completely supressed in the active droplets. Intriguingly, some particles formed from the short peptide and active ribozyme system initially adopted a non-spherical morphology and relaxed to form round droplets over the course of the experiment **(Figure 2 – supplement 4)**, similar to morphological changes induced by the extension of UDP in spermine-based coacervates^32^. This indicates that the material properties of the droplets change over time.

**Table 1.**
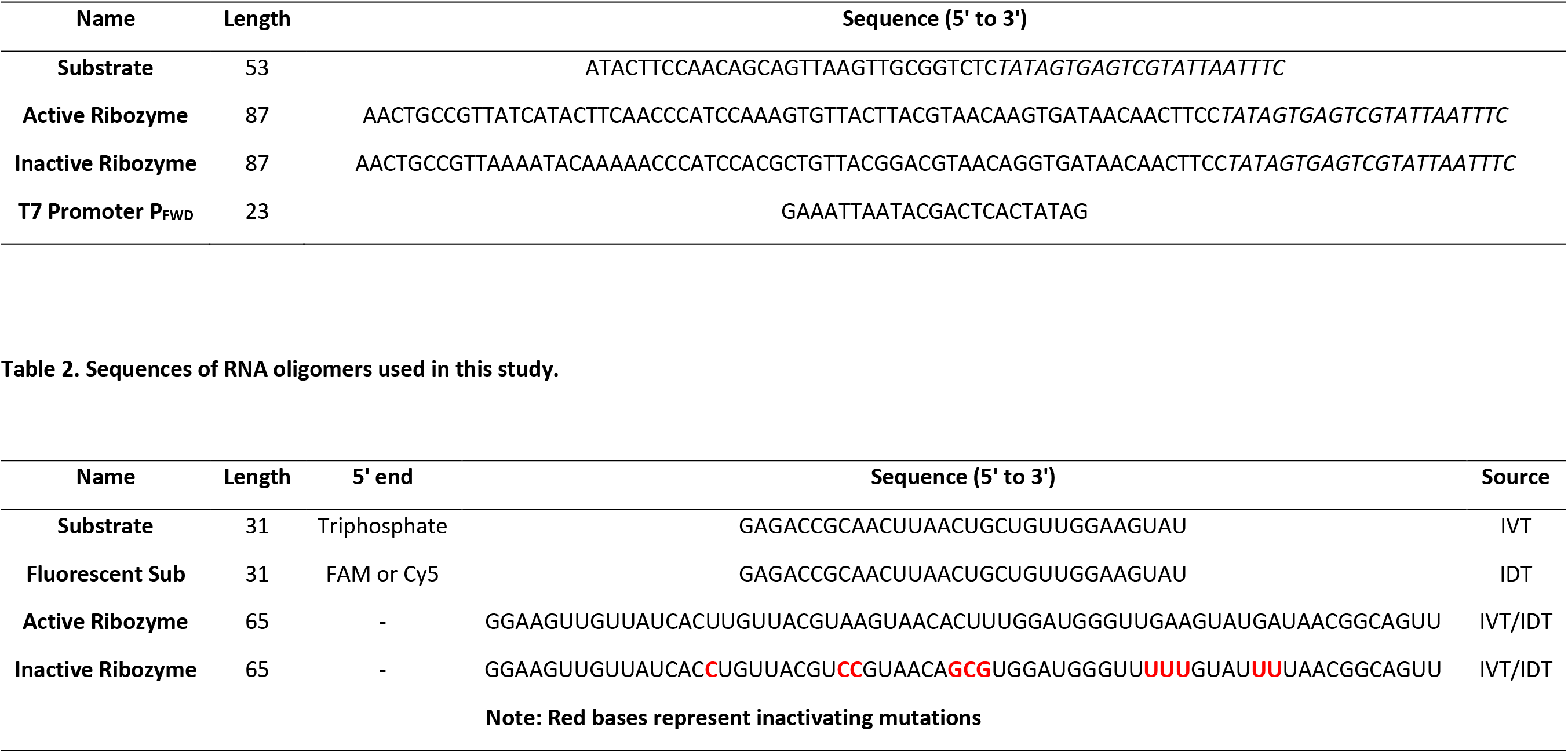
Sequences of DNA oligomers used in this study.

**Table 2.** Sequences of RNA oligomers used in this study.

**Figure 2.**
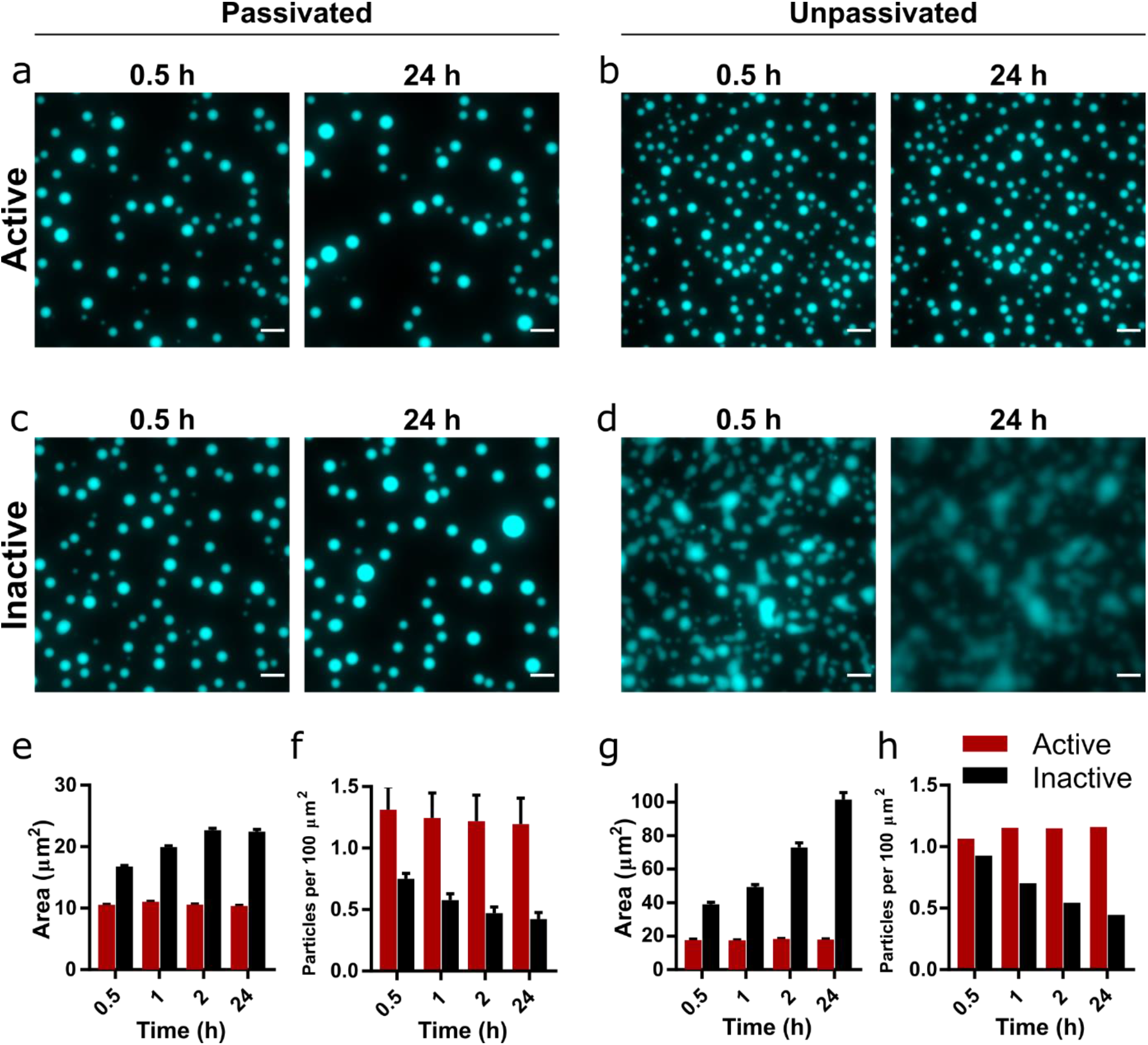
Droplet morphology over time in active and inactive coacervate systems formed from R3C RNA and (Lys)_19-72_. Images of coacervate droplets prepared with active **(a, b)** or inactive ribozyme **(c, d)** and 0.75:1 (Lys)_19-72_:RNA in passivated **(a, c)** and unpassivated **(b, d)** environments. Scale bars = 10 μm. Average particle areas and population density over time for the passivated environment are shown in **e** and **f** respectively. Equivalent data for the unpassivated environment are shown in **g** and **h**. All experiments were performed with 1 mM total RNA monomer concentration, a 0.75:1 ratio of (Lys)_19-72_:RNA monomers, and at 30 °C, pH 8.6 and 10 mM MgCl_2_. The RNA reaction mixture contained 10 % Cy5-labelled substrate for fluorescence imaging. Particles were measured from at least 9 separate images, except for unpassivated samples for which a single image was captured. Error bars are standard errors. Droplet areas and particle counts were measured using the CellPose segmentation algorithm. Data for condensates formed from the shorter (Lys)_5-24_ peptide are shown in **Figure 2 – supplement 2**.

Whilst surface passivation is used as standard in the imaging of coacervate systems to prevent wetting and adhesion effects, surface interactions can affect droplet formation and morphology, and therefore may also influence the observed protocellular phenotype. Consequently, we repeated the previous experiment on an unpassivated polystyrene surface (Greiner μclear microplate, medium binding). The coacervate droplets containing the active ribozyme behaved as previously observed, with discrete round morphologies **(Figure 2b)** and little change in both average droplet area and population density over the course of the experiment **(Figure 2g, h)**. These droplets exhibited greater average areas than those in the passivated environment, likely due to wetting onto the surface. In contrast, droplets containing the inactive ribozyme wet the surface and rapidly spread, eventually merging to form a film of the condensed coacervate phase on the bottom of the well **(Figure 2d)**. The measured average particle area therefore increased greatly over the course of the experiment, whilst the number of particles decreased **(Figure 2g, h)**.

These differences in behaviour indicate that the generation of longer RNA within the coacervate environment imparts altered physical properties on the droplets, despite maintaining a spherical morphology typical of a liquid system. To further investigate this phenomenon, we added the (Lys)_5-24_ and (Lys)_19-72_ peptides to pre-reacted RNA mixtures. Here, we observed that mixtures containing the active ribozyme and therefore pre-concatenated RNA initially formed non-spherical gel-like condensates with both peptides, which relaxed to form spherical droplets over the course of 24 h, whilst mixtures with inactive ribozyme yielded spherical droplets from the outset (**Figure 2 – supplement 5**). This suggests that large morphological differences between active and inactive systems are not observed when peptide is added to ribozyme and substrate mixtures without pre-reaction because the average RNA length is initially identical. As the reaction proceeds in the active systems, a transition to a more viscous or gel-like state may occur whilst maintaining the initially formed spherical morphology.

Given the effect of RNA concatenation on the coacervate droplets, we asked whether the activity of the E_L_ ribozyme could affect the interactions between different populations of droplets. Using the previously established conditions and concentration ratios, we mixed populations of droplets containing either a Cy5- or FAM-tagged substrate (10 % total substrate concentration) and monitored mixing and content exchange over the course of 24 h. These effects can be quantified by the calculation of a Pearson correlation coefficient (PCC), which measures the correlation of pixel intensities between the two fluorescence channels^44^. Two coefficients were calculated: PCC_droplet_, which measures the degree to which fluorophores mix within individual fused droplets, and PCC_pop_, which is calculated on a population level and measures the degree of mixing between the two populations. Positive values indicate spatial colocalization of fluorophores, whilst negative values indicate spatial separation of fluorophores and therefore the presence of discrete regions or populations.

Shortly following mixing, discrete populations of Cy5- and FAM-labelled droplets are clearly visible **(Figure 3a, b and Figure 3 – supplement 1)**. Fused and unevenly mixed droplets containing both fluorophores are visible, providing evidence that coalescence is the mechanism of droplet growth in this system. After 24 hours, visual inspection of the images reveals that discrete populations of Cy5- and FAM-labelled droplets only persist in droplets containing the active ribozyme **(Figure 3a and Figure 3 – supplement 1a)**. In the inactive droplets, both fluorophores appear evenly distributed throughout the population **(Figure 3b and Figure 3 – supplement 1b)**. In all cases, the PCC_droplet_ increased over time, tending towards unity, indicating that labelled RNA was able to equilibrate within the droplets via diffusion **(Figure 3c and Figure 3 – supplement 1c)**. However, it is notable that the rate and magnitude of increase in PCC_droplet_ was greater in droplets containing the inactive ribozyme (**Figure 3 – supplement 2**), indicating greater RNA mobility. For active and inactive (Lys)_19-72_ droplets, PCC_pop_ was initially negative, implying discrete and distinct populations **(Figure 3d)**. Over the course of the experiment, the active system maintained this separation, whilst the PCC_pop_ of the inactive system tended towards unity. These trends are further confirmed by scatter plots of normalized FAM and Cy5 fluorescence intensity in individual droplets (**Figure 3 – supplement 3**), which show the clearly separated populations of active systems and mixing in inactive systems. As observed in our previous experiments, mixtures of the active ribozyme and peptide produced smaller droplets which did not grow over the course of the experiment compared to inactive populations, suggesting that droplets with inactive ribozyme are more prone to coalesce and therefore mixing than droplets containing the active ribozyme **(Figure 3e, f)**.

**Figure 3.**
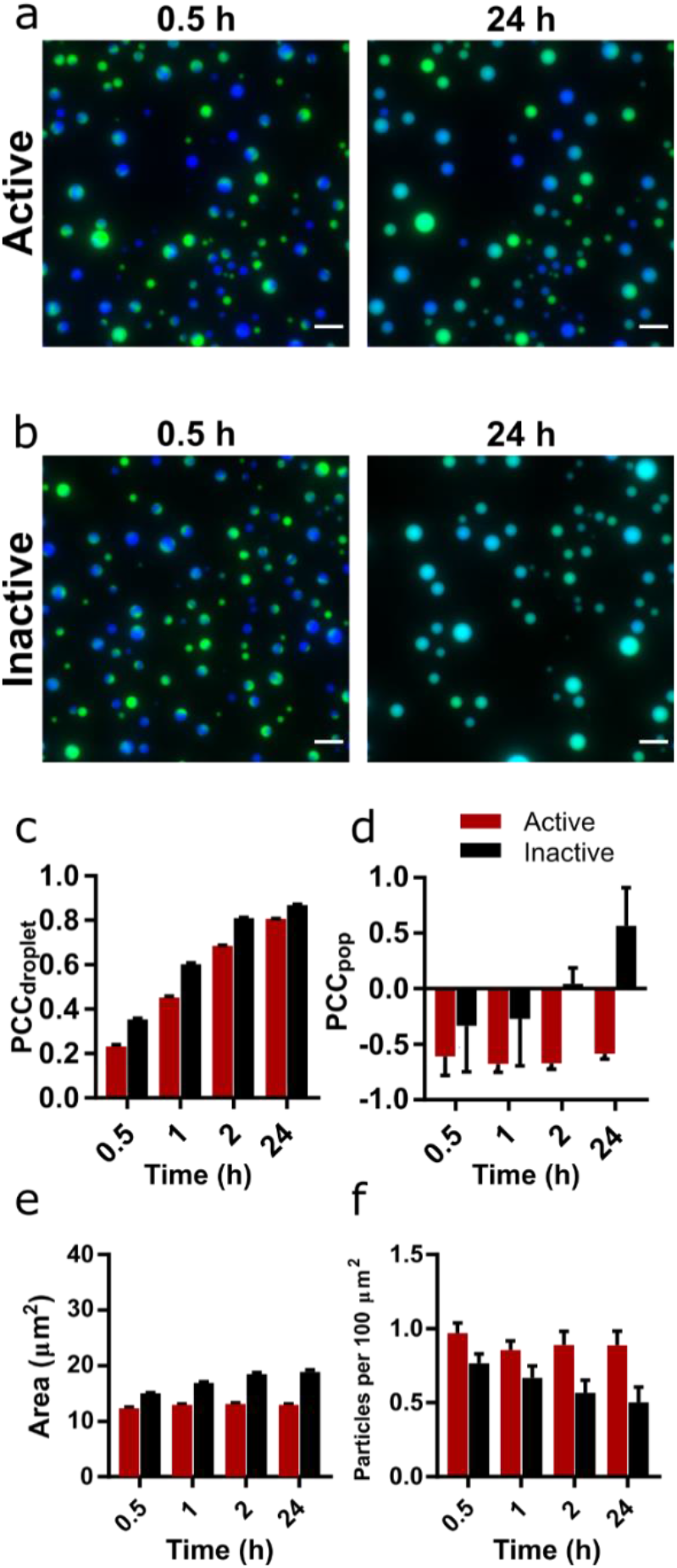
Mixing and content exchange between populations of active or inactive coacervate systems formed from R3C RNA and (Lys)_19-72_ peptide. **a)** and **b)** Example images of mixtures of orthogonally labelled coacervate droplets prepared with 0.75:1 (Lys)_19-72_:RNA containing either active (**a**) or inactive (**b**). Each population in the set contained either 10 % FAM- or 10 % Cy5-tagged substrate (green and blue, respectively). The two populations in each set were mixed shortly after preparation and then imaged over 24 h in a passivated environment. All experiments were performed at 30 °C, pH 8.6 and 10 mM MgCl_2_ with a 1 mM total RNA monomer concentration and a 0.75:1 ratio of (Lys)_19-72_:RNA. The colocalization of the two fluorophores within single droplets is measured by the droplet Pearson coefficient (PCCdroplet) **(c)**, whilst the colocalization of fluorophores in the overall population of droplets is measured by the population Pearson coefficient (PCCpop) **(d)**. The average particle area and number of particles per unit area over time are shown in **e** and **f** respectively. Particles were measured from at least 6 separate images. Error bars are standard errors. Scale bars = 10 μm. Data for condensates formed from the shorter (Lys)_5-24_ peptide are shown in **Figure 3 - supplement 1**.

A similar yet weaker trend was observed with short (Lys)_5-24_ peptide droplets, which exhibited greater content exchange overall between both active and inactive droplets **(Figure 3 – supplement 1–3)**. Here, the presence of the active ribozyme slowed the rate of coalescence and therefore content exchange rather than completely suppressing it, but still allowed active populations to maintain a limited degree of identity over the course of the experiment.

## Discussion

Our original goal was to develop a ribozyme capable of RNA chaining that could significantly increase the average RNA length in a coacervate system, and thereby allow us to study the influence of such a reaction on the material properties of the coacervates. Although ribozyme systems that concatenate a short substrate have been previously reported, these do so via a reversible cyclic-phosphate mediated mechanism, and as such the concentration of extended products decreases rapidly as product length increases^45^. The R3C ligase ribozyme was thus an attractive starting point for the development of our system as the 5’-triphosphate activated reaction it catalyzes is quasi-irreversible, and the ribozyme is already optimized for high rate and yield^39^. Indeed, the E_L_ ribozyme was able to generate RNA strands of > 500 nt in length, including circular species, and this activity was enhanced in terms of both rate and yield in (Lys)_n_ coacervates. Although the observation of circular products in concatenation reactions has been previously reported in ribozyme systems^46^, the inhibition of this behaviour by coacervation is notable. Whilst the mechanism of this inhibition is unknown, the absence of circular products suggests that the ends of a substrate strand do not meet, hinting at a decrease in RNA mobility, increased chain stiffness or reduced release of ligated substrate from the ribozyme within the condensed phase compared to the bulk solution. The suppression of circularisation should lead to higher overall RNA lengths as the concentration of RNA ends available for ligation is not decreased over the course of the reaction. It is not possible to quantify this difference as measurement of the average chain lengths requires the use of a 5’ fluorescent tag, which itself prevents formation of circular products due to the blocked 5’ end.

We have previously reported that the activity of the hairpin ribozyme (HPz) is greatly enhanced in condensed phases comprised of catalytic RNA and (Lys)_n_ peptides^15^. The increase in turbidity observed on titrating either (Lys)_19-72_ or (Lys)_5-24_ into a fixed concentration of R3C ribozyme and substrate was similar to that observed for HPz, although in the present study the resultant condensed phase appeared liquid rather than gel-like. This variation in condensed phase morphology may be due to the influence of RNA structure, sequence and hybridisation state on condensation^47,48^, with a higher degree of single stranded or loop structures in the E_L_ ribozyme system. The inhibition of ribozyme activity at excess (Lys)_19-72_:RNA ratios was comparable to that observed in our previous study^15^, and is attributed to a peptide length dependent melting or misfolding of native nucleic acid tertiary structures at high concentrations of longer peptides^49,50^. Our previous study into the function of the R3C replicase ribozyme in RNA-peptide coacervates reported a similar trend: catalytic activity was inhibited at excess (Lys)_n_:RNA ratios for a range of peptides (n = 7, 8, 9, 10, 18, and 19-72)^14^. The present study extends the peptide concentration range in which the R3C ribozyme functions, with robust catalytic activity at ratios as high as 4:1 (Lys)_5-24_:RNA. A possible explanation is that commercially available (Lys)_n_ deviates from the manufacturer’s stated size range, with the (Lys)_5-24_ used here being predominantly comprised of oligomers between n = 3 to n = 9 in length^15^. These short fragments may interact only weakly with the RNA, effectively reducing the concentration of peptide oligomers that can form condensates. In general, it should be noted that the effect of phase separation on ribozyme activity in a given system is highly specific to the identity of the ribozyme and peptide, their relative concentrations, and environmental conditions such as buffer, magnesium concentration and temperature.

The difference in growth behaviour between active and inactive droplets is a striking demonstration of how the presence of a catalytic species can modulate the physical properties and behaviour of a model protocellular system, in this case by reducing coalescence. The coarsening of liquid coacervate phases is not inevitable: Growth can be also be suppressed by active chemical processes hosted within droplets that produce the polymer components^51^, although in our experiments the action of the ribozyme does not increase the concentration of RNA or peptide in the system. The formation of kinetically trapped states can also allow the persistence of static droplet populations over time^52^. However, increasing polymer lengths in a coacervate system results in stronger cooperative electrostatic interactions between polymer components^34^, and has been shown to affect physical parameters such as droplet water content, critical salt concentration and condensed phase polymer concentration^35^, leading to denser droplets and a more depleted dilute phase. Simulations have shown that increasing RNA length modulates both the material and interfacial properties of RNA – peptide condensates, in particular increasing density and surface tension^53^. RNA length also has been shown to modulate viscosity in biomolecular condensates^54^, which in turn determines the droplet fusion dynamics^55^. Thus, increasing overall RNA length by R3C substrate concatenation may inhibit growth via coalescence by increasing condensate density and viscosity, and reduce wetting affects through the alteration of surface tension. Whilst RNA sequence and secondary structure have also been shown to influence the viscoelastic behaviour of RNA peptide condensates^56^, the active and inactive ribozyme systems used in the present study differ by only few nucleotides.

Despite the spherical morphology of the RNA – peptide condensates reported here, the inhibition of growth and wetting may also be due to a phase transition to a gel state. The formation of non-liquid RNA-peptide condensates has been previously reported for poly-rA:rU-peptide mixtures, where RNA base pairing leads to the formation of a kinetically arrested gel-like solid^48^, as well as in our previous work^15^. These gel condensates melt to form spherical particles upon thermal denaturation and annealing, which disrupts the networked structures formed from complementary RNA strands and peptides. In our R3C system, complementary base-pairing interactions between substrate and ribozyme should also permit the formation of large, non-covalently assembled networked structures due to the repeating nature of the substrate. Ribozyme activity is expected to increase the stability of these structures, as substrate ligation increases the free energy of association between ribozyme and substrate. We hypothesise that in the inactive system, and before significant concatenation occurs in the active system, RNA-RNA interactions are sufficiently weak to allow the formation of liquid coacervate droplets. In the active system, liquid droplets initially form because the timescale of coalescence is faster than that of RNA ligation. However, as ligation proceeds the stability of networked assemblies of ribozyme and substrate increases, resulting in a transition from a liquid to either a highly viscous liquid state or solid gel state that is both unable to grow and or wet the unpassivated surface on the timescale of the experiment. This hypothesis is supported by our observation that mixtures of pre-concatenated RNA and (Lys)_n_ initially form non – spherical gel-like aggregates, which then relax to form spherical droplets over the course of 24 h.

The reduction of mixing between populations of droplets containing orthogonally labelled RNA likely results from the changes in material properties and growth behaviour caused by RNA concatenation. The coalescence of the coacervate droplets observed in all inactive systems is likely to be the primary mechanism of mixing in this scenario. Indeed, greater droplet growth was observed in the two-population system than for single population experiments, which may be due to the necessity of pipetting each droplet population into the sample environment sequentially, thus increasing mechanical agitation and contacts between droplets. All populations at the initial timepoint contain fusion droplets, which contain separate and unmixed areas of each fluorescent RNA. Notably, fusion droplets in all systems appear evenly mixed after 24 h, suggesting that significant equilibration occurs before any transition to a highly viscous liquid or solid gel. Mixing may also occur by diffusive transfer of RNA between droplets. As the fluorescently labelled substrate is ligated onto the growing concatemer chain, the diffusion coefficient of the labelled material is expected to decrease over the course of the reaction as its effective length increases, which would in turn slow diffusive transfer in the active system.

Taken together, the results reported here describe a range of altered droplet behaviours stemming from ribozyme-driven changes in coacervate physical properties. The ability of a population of protocells to resist passive coarsening, surface wetting and content exchange can be considered a fitness advantage^57^, and the emergence of fitness differences from varying protocellular compositions and phenotypes is a key step on the path towards Darwinian evolution with suitable selective pressures^58^. Furthermore, in this system the phenotypic difference originates from the ribozyme-driven concatenation reaction, which increases the length of the anionic coacervate component. The sequence of the ribozyme in effect comprises the genotype of our protocellular system, with different variants (e.g. active or inactive sequences) producing different protocellular phenotypes. The realisation of a phenotype-genotype linkage is essential for true open-ended evolution, otherwise phenotypes with obvious fitness advantages have no way of being propagated to future generations. We envision that recent methodological developments in RNA sequencing from single coacervate microdroplets will enable future selection experiments on populations of protocells with varying RNA genotypes and varying degrees of fitness with respect to environmental pressures such as heat or salt concentration^59^. It should be noted that as a ribozyme developed by *in vitro* selection, the R3C ligase is already a highly optimized system and offers little potential for further enhancement via selection. Nevertheless, the reciprocal modulation of ribozyme activity and coacervate properties reported here furthers the argument for the early coevolution of RNA and peptides^7–9,18,60^. Whilst the magnitude of behavioural change was greater for active and inactive coacervate populations formed with the longer peptide, clear phenotypic differences were nonetheless also present in the (Lys)_5-24_ system. This short peptide, predominantly composed of n = 3 – 9 residue oligomers, is of a length that could be produced by prebiotically plausible processes such as wet-dry cycling^61,62^. Although the simple lysine-based peptides used here are a basic model system, additional variation in droplet properties may be achieved in future by varying peptide sequence. Droplet populations formed from different peptides could exhibit varying degrees of ribozyme enhancement, or fundamentally different physical properties that allow selection based on peptide identity as well as RNA sequence and activity.

## Materials

Trizma base (Tris) (Thermo Fisher Scientific, Waltham, MA), sodium hexametaphosphate ((NaPO_3_)_6_, 611.77 g/mol) (Sigma-Aldrich, St. Louis, MI), formamide (CH_3_NO, 45.04 g/mol) (Sigma-Aldrich), ethylenediaminetetracetic acid disodium salt dihydrate (EDTA, C_10_H_14_N_2_Na_2_O_8_·2H_2_O, 372.24 g/mol) (Sigma-Aldrich), magnesium chloride hexahydrate (MgCl_2_·6H_2_O, 203.30 g/mol) (Merck, Darmstadt, Germany), sodium hydroxide (NaOH, 39.997 g/mol) (VWR, Radnor, PA), sodium chloride (NaCl, 58.44 g/mol) (Sigma-Aldrich), urea (CH_4_N_2_O, 60.06 g/mol) (Carl Roth, Karlsruhe, Germany), hydrochloric acid (HCl, 37 %, 36.46 g/mol) (VWR), boric acid (H_3_BO_3_, 61.83 g/mol) (Merck), ammonium persulfate (APS, (NH_4_)_2_S_2_O_8_, 228.20 g/mol) (VWR), acrylamide (19:1 bisacrylamide) (Thermo Fisher Scientific), tetramethylethylendiamine (TEMED, C_6_H_16_N_2_, 116.21 g/mol) (Carl Roth), SYBR gold stain (Thermo Fisher Scientific), RNA oligomer length standard (Low range ssRNA ladder) (NEBm Ipswich, MA).

All peptides were purchased from Sigma-Aldrich and used without further purification (poly-L-lysine hydrobromide (4–15 kDa, 19-72 residues, monomer: 209 g/mol), poly-L-lysine hydrobromide (1-5 kDa, 5-24 residues, monomer: 209 g/mol). All RNA and DNA oligomers were obtained from Integrated DNA Technologies (Coralville, IA) and are listed in Tables 1 and 2. Fluorescently labelled oligomers were ordered with HPLC purification, whilst unlabelled oligomers were desalted and used without further purification. All RNA oligomers were dissolved in RNAse-free water and stored at −80 °C.

## Methods

### Preparation of RNA

RNA was ordered from IDT or transcribed in-house from DNA templates. DNA sequences containing a T7 promoter upstream of the RNA sequence of interest were ordered from IDT. The DNA templates were prepared by annealing a complementary oligo to the T7 promoter region by heating equimolar mixture of the oligos at 85 °C then cooling on ice. The resulting partially double stranded DNA was used as DNA template for transcriptions. Large scale *in vitro* transcription (IVT) reactions were adopted to produce enough RNA for downstream applications. The transcription reaction volume varied from 400 μL – 4 mL and contained the following: 1 μM partially double-stranded DNA template, 30 mM Tris-HCl pH 7.8, 30 mM MgCl_2_, 10 mM DTT, 2 mM Spermidine, 5 mM of each NTP, 1 U/mL Inorganic Pyrophosphatase, 0.5 μM T7 RNA polymerase (purified from a recombinant source in-house). The reaction proceeded for 4-6 hours at 37 °C, after which the volume was concentrated by spinning at 15,000 × g, 4 °C in Amicon ultrafiltration columns (Merck) with 3 kDa molecular weight cut-off regenerated cellulose filters. Following concentration, the RNA was purified with the Monarch RNA cleanup kit (NEB) following manufacturer’s instructions, eluted twice in water and quantified on the NanoDrop OneC (Thermo Fischer Scientific). The resulting ribozymes were gel purified using 12 % urea PAGE, whereas the substrate was purified using 20 % PAGE. The amount of RNA loaded per well was 5 μg mm^−2^ of the area of the bottom of the well. After PAGE, the gel was wrapped in plastic foil and the band of interest was identified by UV shadowing (254 nm, < 30 seconds) on an autofluorescent background. The gel slice was excised, crushed, weighed and soaked in 2 μL of 0.3 M sodium acetate (pH 5.2) per mg of gel at 4 °C overnight. Following elution, the gel debris was removed using Costar Spin-X columns (Corning Inc., Corning, NY) with 0.45 μM cellulose acetate filters. Next, 20 ng RNA - grade Glycogen (Invitrogen, Waltham, MA) and 1.2 volumes of cold isopropanol were added to the solution. The mixture was cooled for 1 hour at −20 °C to promote precipitation, and then centrifuged for an additional hour at 21,000 × g, 4 °C. The supernatant was removed, and the pellet washed twice with 0.5 volumes of cold 80 % ethanol. After washing, the supernatant was completely removed, the, the pellet was dried for a few minutes under vacuum and resuspended in ultrapure water. RNA was aliquoted and stored at −80 °C.

### Urea Polyacrylamide Gel Electrophoresis (PAGE)

Rotiphorese Gel 40 (19:1) was used to prepare 20 % polyacrylamide gel stocks containing 8 M urea and 1X TBE (89 mM Tris, 89 mM Boric Acid, 2 mM EDTA, pH 8). A 0 % gel stock was prepared using the same volumes with water replacing acrylamide. The two stocks were mixed at different ratios to obtain the final desired acrylamide concentration. Polymerization was initiated by adding 0.01 volumes of 10 % APS and 0.001 volumes of TEMED. The gel was cast in an EasyPhor PAGE Maxi Wave (20 cm × 20 cm) (Biozym, Hessisch Oldendorf) and allowed to polymerize at room temperature for > 2 hours, then pre-run at a constant power of 20 W for 45 minutes. The gel thickness was 2 mm and 1 mm for preparative and analytical gels, respectively. The quenched samples were loaded, and the gel was run for 90 minutes at 20 W constant power in 1X TBE running buffer. When necessary, the gel was removed from the glass plates and stained with SYBR Gold (Invitrogen) nucleic acid staining dye for 10 minutes in 1X TBE and 1X SYBR Gold, then washed twice for 5 minutes in de-ionized water to decrease background fluorescence. Images were acquired using and Azure Sapphire RGB gel scanner (λ_ex_ = 520 nm for SYBR Gold, 658 nm for Cy5) and analyzed with the AzureSpot software (Azure Biosystems, Dublin, CA). The images were background subtracted using the built-in rolling ball function set to a diameter of 1000.

### Poly-L-lysine titration PAGE

Ribozyme assays were carried out with a 10 μL total reaction volume and the following components: 50 mM Tris-HCl pH 8.6, 10.5 μM ribozyme, 10.5 μM substrate, 10 mM MgCl_2_ and varying concentrations of poly-L-lysine. Ratios of RNA (Lys)_n_ were (based on the charge concentration of the RNA (1 mM), calculated with the following equation:

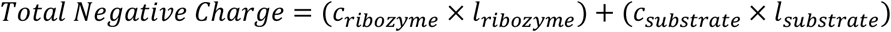

Positive charge concentrations were calculated based on the lysine hydrobromide monomer repeat molecular weight (209 g/mol) and mass of poly-L-lysine, considering a single positive charge for each residue. Reactions were set up at room temperature by first adding the all the components except the RNA to allow equilibration of poly-L-lysine in the buffer. The reaction was then started by adding a mixture of ribozyme and substrate and incubated in a thermocycler at 30 °C for 2 hours. Reactions were stopped by adding 1 volume of 5 M NaCl, 1 volume of 1.25 M hexametaphosphate (HMP) and 12 volumes of RNA loading buffer containing 10 mM EDTA, 0.05 % bromophenol blue, 95 % Formamide. The resulting samples were briefly vortexed, denatured for 5 minutes at 85 °C, cooled quickly on ice and centrifuged for 5 minutes at 2000 x g (Color Sprout Plus, Biozym). PAGE analysis proceeded as described above.

### OD measurements

Measurements were performed using a NanoDrop OneC (Thermo Fischer Scientific) by measuring absorbance at 500 nm. The RNAs were pre-reacted at 2X concentration (21 μM each) in 1X buffer (50 mM Tris-HCl pH 8.6, 10 mM MgCl_2_) for 3 hours at 30 °C. Varying concentrations of poly-L-lysine in 1X buffer were then added to the reacted RNA. For every poly-L-lysine concentration tested, 2 μL of pre-reacted RNA was added to 2 μL of poly-L-lysine and mixed by pipetting 10 times. The resulting solution was incubated for 5 minutes, after which absorbance was measured. At least 3 biological replicates were measured for each concentration. From these data, we selected specific peptide:RNA ratios for both peptides for further experimentation (0.75:1 (Lys)_19-72_):RNA and 3:1 (Lys)_5-24_:RNA). The criterion for this selection was the formation of liquid coacervate droplets at a peptide:RNA ratio that did not supress ribozyme activity. Both concentrations occur shortly before the respective turbidity maxima in the peptide titration turbidity curve.

### Ribozyme kinetics

Time-dependent assays were performed using individual aliquots for each time point to mitigate the effects of coacervate adhesion to the PCR tube walls over time. The reaction mixture contained: 50 mM Tris-HCl pH 8.6, 10 mM MgCl_2_, 1 mM RNA charge concentration (9.5 μM substrate, 1 μM Cy5-tagged substrate, 10.5 μM ribozyme), and either 3 mM (Lys)_5-24_ or 0.75 mM (Lys)_19-72_ charge concentration. Aliquoted reactions were quenched by the addition of 2 μL 5 M NaCl, 2 μL 1.25 M HMP, and 34 μL RNA loading buffer (10 mM ETDA, 0.05 % bromophenol blue, 95 % formamide) at designated time points (t = 0, 2, 5, 10, 20, 30, 60, 120, 180, 240 minutes). Sample preparation and PAGE were performed as described above. The average substrate length and relative band percentages were calculated as follows:

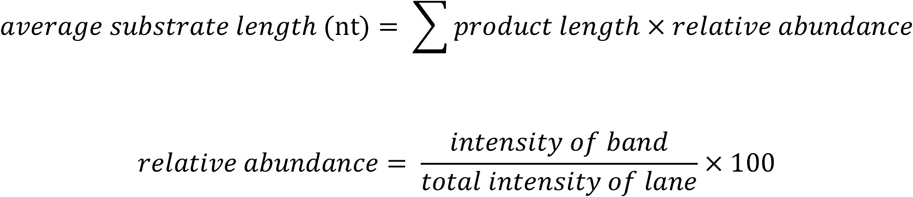

The data were fitted in GraphPad Prism using both a first order (‘one-phase association’) and second order (‘two-phase association’) kinetic model. First and second order kinetics were discriminated between using the extra sum-of-squares F test for nested models. The simpler first order model was rejected when P < 0.05.

### Temperature and Ratio Screen

Reactions were carried out with varying ribozyme to substrate ratios, maintaining a total negative charge of 1 mM. Reactions with component ratios of 1:1, 1:2, and 1:4 contained ribozyme:substrate concentrations 10.5:10.5 μM, 8:16 μM, and 5:20 μM respectively. Buffer conditions were 50 mM Tris-HCl, pH 8.6, and 10 mM MgCl_2_. The reactions were incubated for 2 hours at 30 °C, 37 °C and 45 °C, then quenched by the addition of 9 volumes of RNA loading buffer (10 mM EDTA, 0.05 % bromophenol blue, 95 % formamide). They were resolved on an 8 % urea PAGE and stained and imaged as described previously.

### RNase R digestion

The reaction products from 10 μL reactions containing 1:1 ribozyme:substrate with either no poly-L-lysine, 0.75:1 (Lys)_19-72-15_ or 3:1 (Lys)_5-24_ and incubated for 3 hours at 45 °C were recovered using a 50 μg Monarch RNA cleanup kit (NEB) and eluted in 12 μL ultrapure water. RNase R (Applied Biological Materials, Richmond, Canada) was used to digest a portion of the purified reaction products. The digest reaction mixture (20 μL) contained 1 μg of RNA, 1X RNase R buffer and 30 U of RNase R, and was incubated for 3.5 hours at 37 °C. The digested RNA was recovered with the 10 μg Monarch RNA cleanup kit (NEB), eluted in 6 μL of ultrapure water and mixed with 6 μL of RNA loading buffer. 1.5 μL of the undigested RNA and 5 μL of the digested RNA were resolved on an 8 % PAGE, stained and imaged as described above.

### Microscopy

Microscopy was performed on a Leica Thunder inverted widefield microscope equipped with a sCMOS camera Leica DFC9000 GTC using a 63x / NA 1.47 objective. Fluorescence channels were λ_Ex_ = 484 - 496 nm / λ_Em_ = 507 - 543 nm for FAM, and λ_Ex_ 629 - 645 nm / λ_Em_ 669 - 741 nm for Cy5. The sample stage was warmed to 30 °C. Samples were loaded into clear bottomed 384 well plates (Greiner μclear, medium binding). Coacervate droplets were formed by combining a solution of RNA with a solution containing poly(L-lysine), buffer and magnesium chloride and mixing using a pipette in a PCR tube. The mixture was incubated on ice for five minutes, then 5 μL was loaded into the well plate. Individual wells were sealed with a drop of silicon oil to prevent evaporation. After loading, plates were immediately incubated at 30 °C. Imaging was performed as soon as the coacervate suspension had settled, typically first at 30 minutes after mixing. A 3 × 3 image grid, centred at the middle of the well, was captured for each sample at various time points. In most cases multiwell plates were passivated using Pluronic F-68 to prevent droplet wetting and adhesion.

### Image processing and analysis

Following data collection grid tiles were inspected, and out of focus images were discarded. Only images that remained in focus for all timepoints in a given sample were carried forward for analysis. Image masks were produced by segmentation in the brightfield channel using the CellPose algorithm (cyto2 model, average cytoplasm diameter = 30 - 100 pixels, flow threshold = 0.4, cell probability threshold = 0)^43^. A rolling ball background was subtracted in all fluorescence channels (radius = 100 pixels). Masks and images were then used to measure the number of particles per image, particle area and volume corrected fluorescence intensity. Both inter- and intra-particle Pearson correlation coefficients were calculated for samples containing two populations of droplets with orthogonal labels. For figure preparation, time course images were aligned using HyperStackReg (v.5.6, translation transformation) and displayed without background subtraction.^63^

## Data availability

All data reported here are supplied within the manuscript. Source Data files have been provided for Figures 1, 2 and 3, including figure supplements where relevant. These include original unedited gel images, microscope images, fitting files, and numeric data. All DNA and RNA sequences are provided in the manuscript.

## Acknowledgements

We wish to thank Martin Spitaler, Markus Oster, Giovanni Cardone and the MPIB Imaging Core Facility for providing excellent guidance throughout the project, as well as subsidized equipment access. H.M. and E.S. were supported by funding from the Deutsche Forschungsgemeinschaft (DFG, German Research Foundation, Project-ID 364653263 – TRR 235). H.M. gratefully acknowledges support by the European Research Council (ERC) under the Horizon 2020 research and innovation program (grant agreement ID: 802000, RiboLife). K.L.V., B.G., T-Y D.T and H.M. were supported by the Volkswagen Foundation with funding from the initiative ‘Life? - A Fresh Scientific Approach to the Basic Principles of Life’ (grant number: 92772).

## Figure supplements

**Figure 1 – supplement 1.**
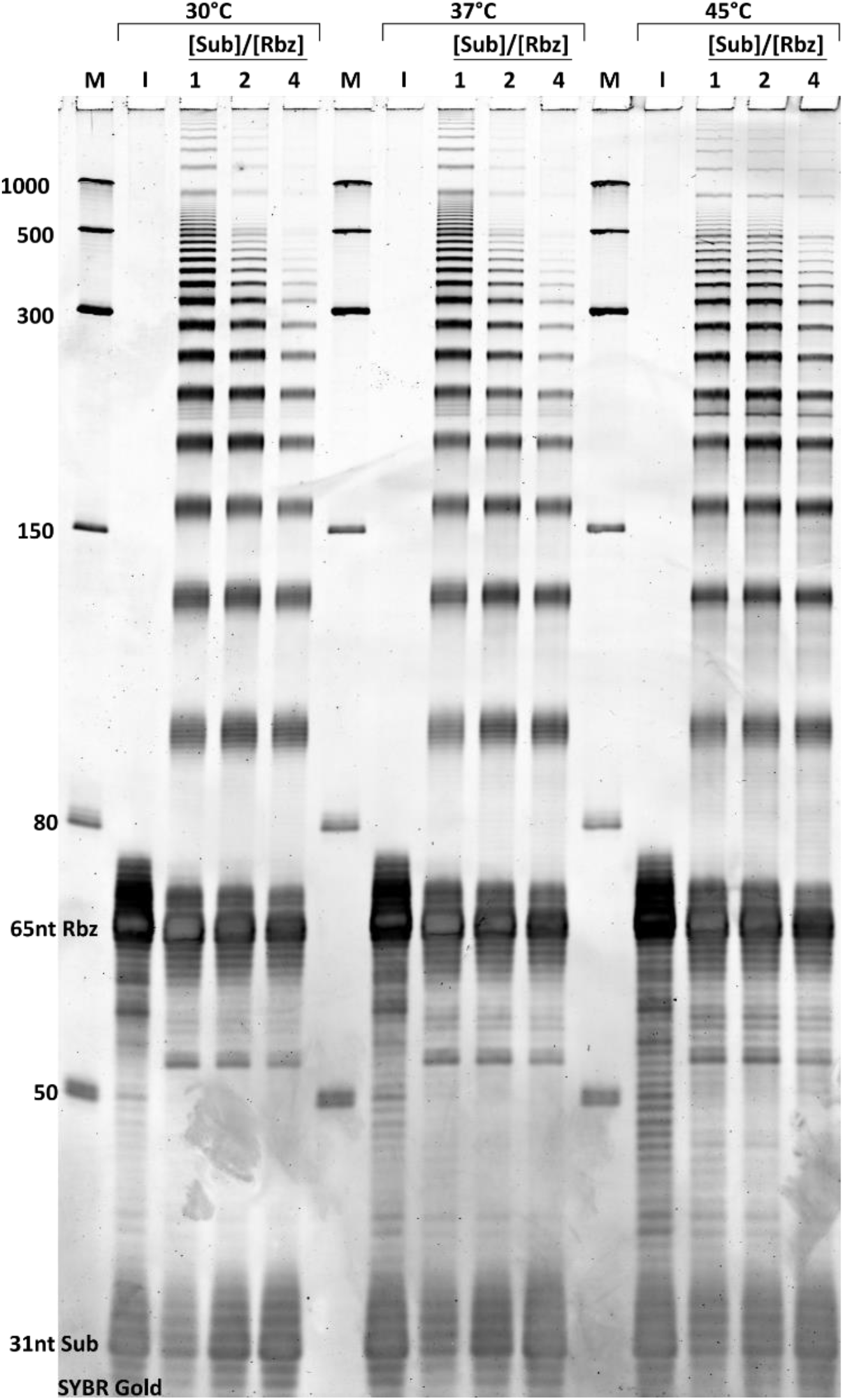
Characterisation of R3C ladder ribozyme activity at various temperatures and ribozyme:substrate ratios. The concatenation activity of the ribozyme was determined by reaction at 30, 37 or 45 ^0^C and with either equimolar, two-fold or four-fold substrate concentration relative to the ribozyme. The reaction buffer contained 10 mM MgCl_2_ and 50 mM tris pH 8.6, and the reaction was stopped after two hours. Excess substrate was found to inhibit the formation of long substrate concatenates at lower temperatures.

**Figure 1 – supplement 2.**
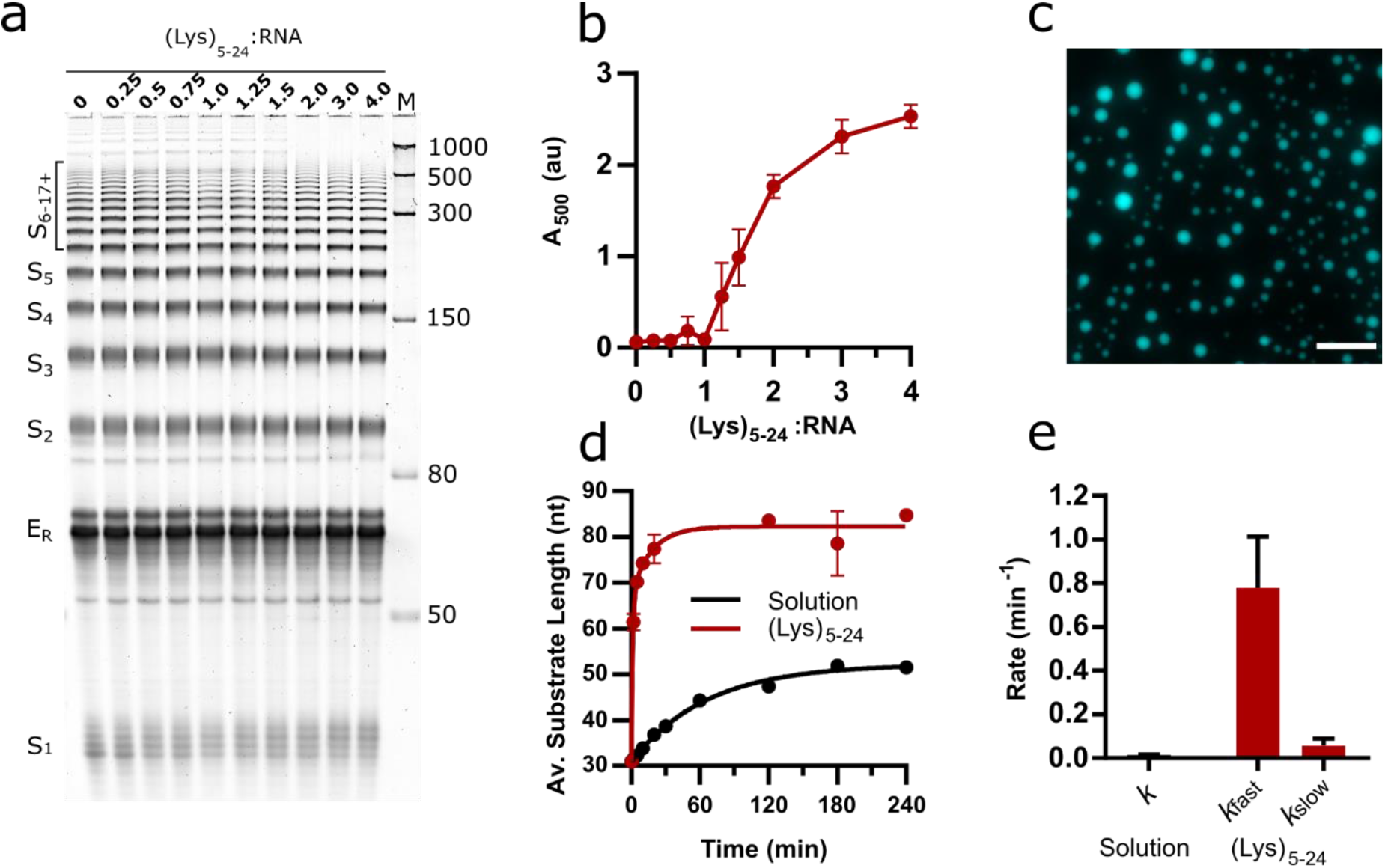
Activity of the E_L_ R3C ladder ribozyme in the presence of short (Lys)_5-24_ peptides. **a)** An 8 % urea PAGE showing the products of the R3C ladder system in solution and varying ratios of (Lys)_5-24_ to R3C RNA (total monomer concentration = 1 mM) after a 2 h reaction at 30 °C, pH 8.6 and 10 mM MgCl_2_. **b)** Variation in absorbance at 500 nm upon addition of varying ratios of (Lys)_5-24_ to the E_L_ RNA before ligation. Data points are an average of three technical replicates with error bars reporting standard deviation. **c)** Example fluorescence microscopy image of (Lys)_5-24_:RNA condensates at a ratio of 3:1 (Lys)_5-24_:RNA, imaged using 10 % Cy5-tagged substrate strand. Scale bars = 20 μm. **d)** Kinetics of chain elongation in solution (black, first order model), and with 3:1 (Lys)_5-24_:RNA (red, second order model) at 30 °C, pH 8.6 and 10 mM MgCl_2_. **e)** Chain extension rate constants for the R3C ladder ribozyme in solution (black, first order model) and in the presence of 3:1 (Lys)_5-24_:RNA (red, second order model). Data points are an average of three technical replicates. Error bars are standard deviations. Poor sample recovery led to an artificially reduced average substrate length at the t = 30 and t = 60 minute timepoints for the reaction in the presence of (Lys)_5-24_. These points were therefore excluded when fitting data.

**Figure 1 – supplement 3.**
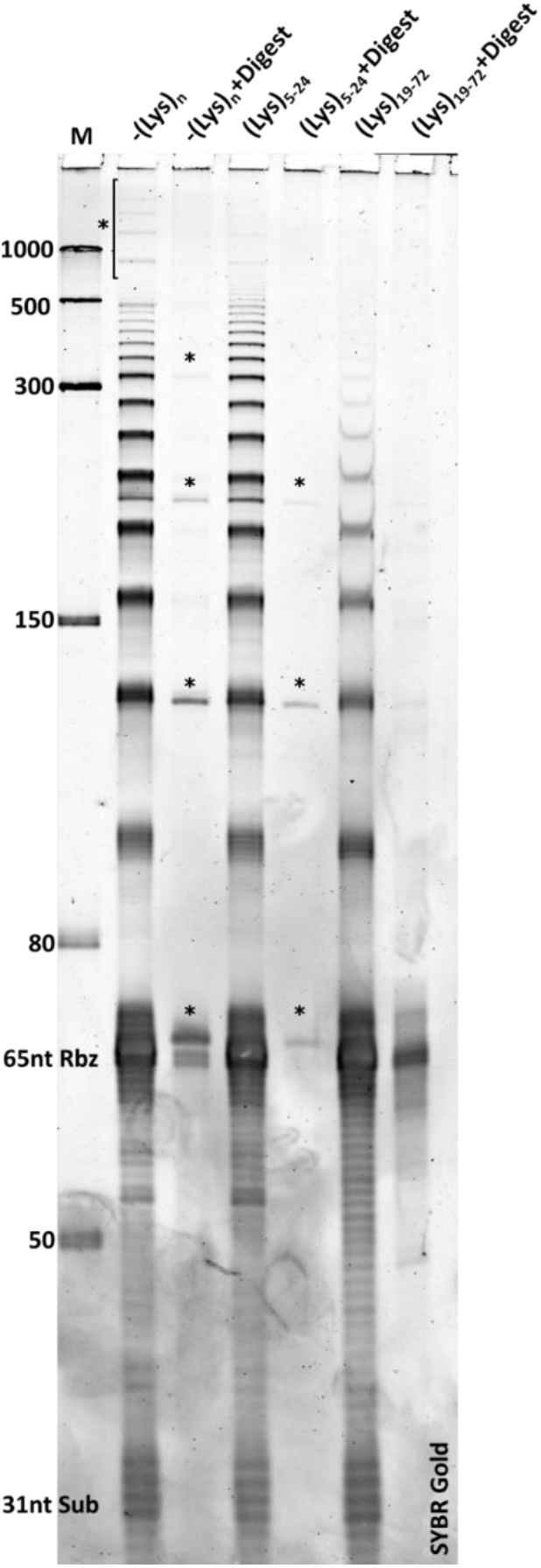
Formation of circular RNA products by the R3C ladder ribozyme. Ribozyme assays were performed at 30 ^0^C in solution and in the presence of poly(lysine) (0.75:1 Lys_19-72_:RNA or 3:1 Lys_5-24_:RNA). The ribozyme reaction buffer contained 10 mM MgCl_2_ and 50 mM tris pH 8.6, and the reaction was stopped after two hours. Circular bands are marked with an asterisk. After two hours the reaction was stopped and the extracted RNA was digested with RNAse R, leaving only circular products. The formation of circular products is observed in solution but is suppressed in the presence of poly(lysine)

**Figure 2 – supplement 1.**
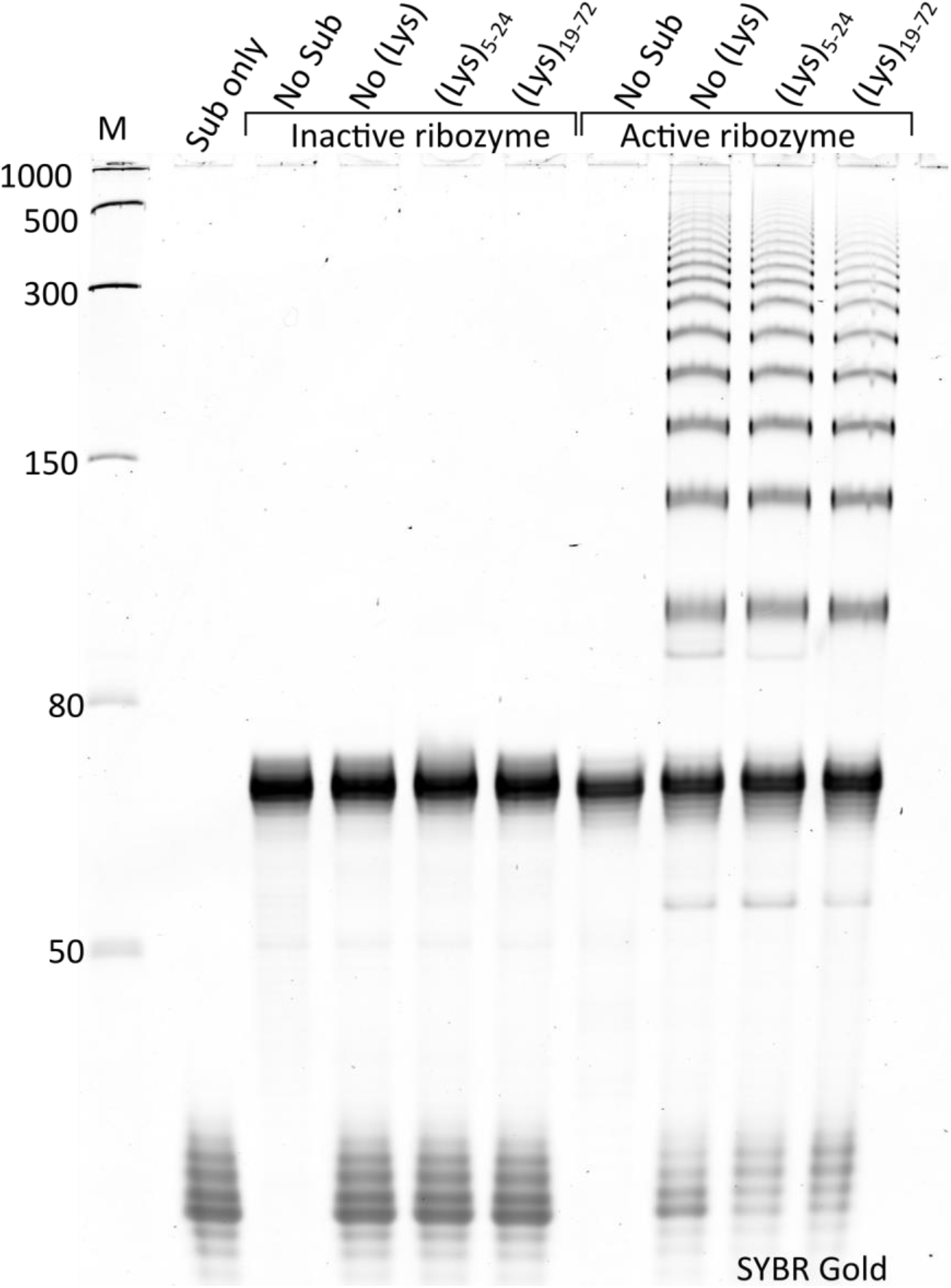
Comparison of active and inactive ribozyme variants. The activity of the active and inactive ribozyme variants was tested in solution and in the presence of poly(lysine) (0.75:1 Lys_19-72_:RNA or 3:1 Lys_5-24_:RNA). The ribozyme reaction buffer contained 10 mM MgCl_2_ and 50 mM tris pH 8.6, and the reaction was stopped after two hours. No ligation activity was detected in the presence of the inactive ribozyme in all conditions tested.

**Figure 2 – supplement 2.**
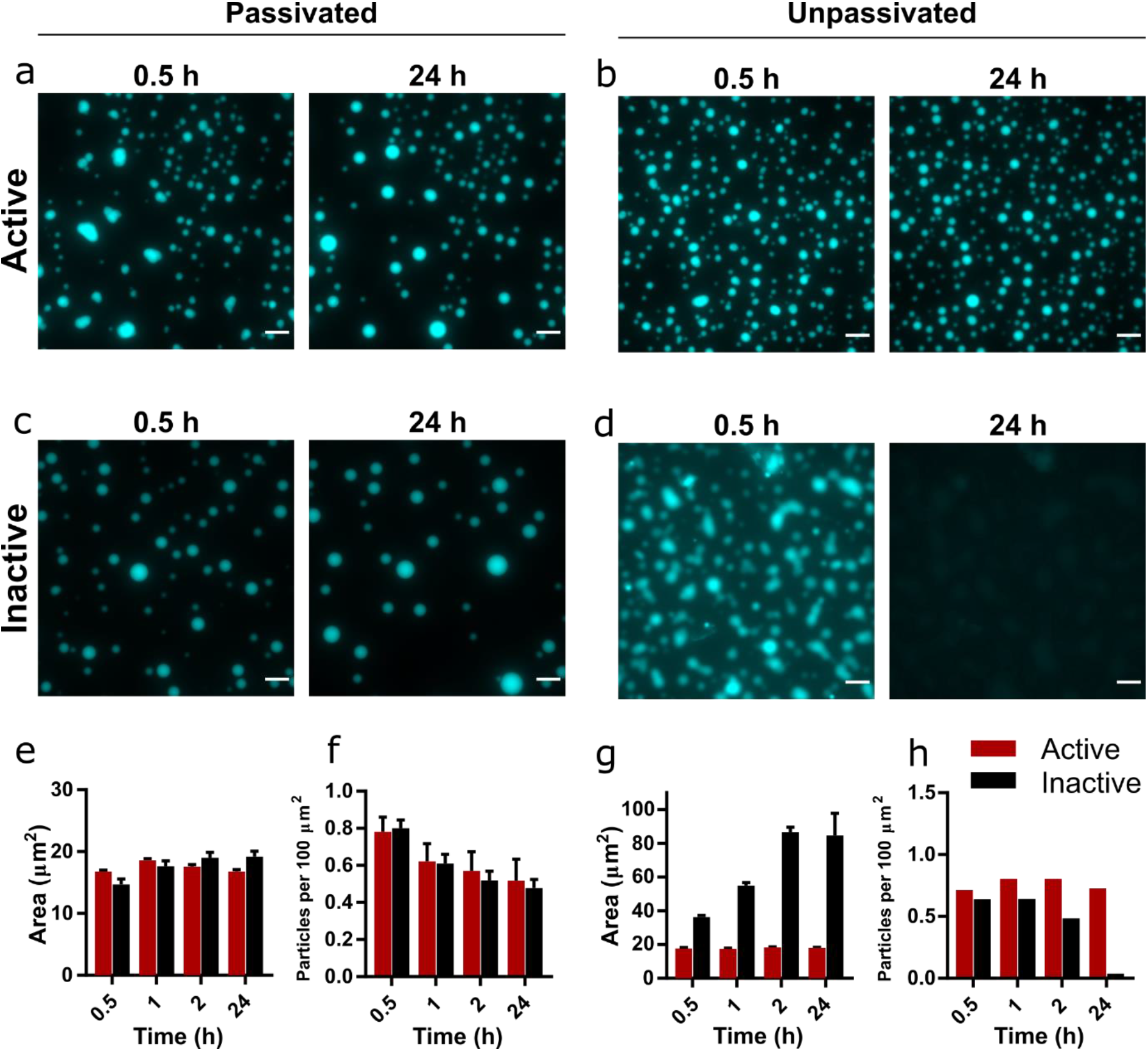
Development of droplet morphology over time in active and inactive coacervate systems formed from R3C RNA and (Lys)_5-24_ peptide. Images of coacervate droplets prepared with active **(a, b)** or inactive ribozyme **(c, d)** and 3:1 (Lys)_5-24_:RNA in passivated **(a, c)** and unpassivated **(b, d)** environments. Scale bars = 10 μm. For the passivated environment, plots of average particle areas and number of particles per unit area over time are shown in **e** and **f** respectively. For the unpassivated environment, these plots are shown in **e** and **f** respectively. All experiments were performed with 1 mM total RNA monomer concentration and a 3:1 ratio of (Lys)_5-24_:RNA. All reactions were performed at 30 °C, pH 8.6 and 10 mM MgCl_2_. Particles were measured from at least 9 separate images, except for unpassivated samples, for which a single image was captured. Error bars are standard errors. The RNA reaction mixture contained 10 % Cy5-labelled substrate for fluorescence imaging. Droplet areas and particle counts were measured using the CellPose segmentation algorithm.

**Figure 2 – supplement 3.**
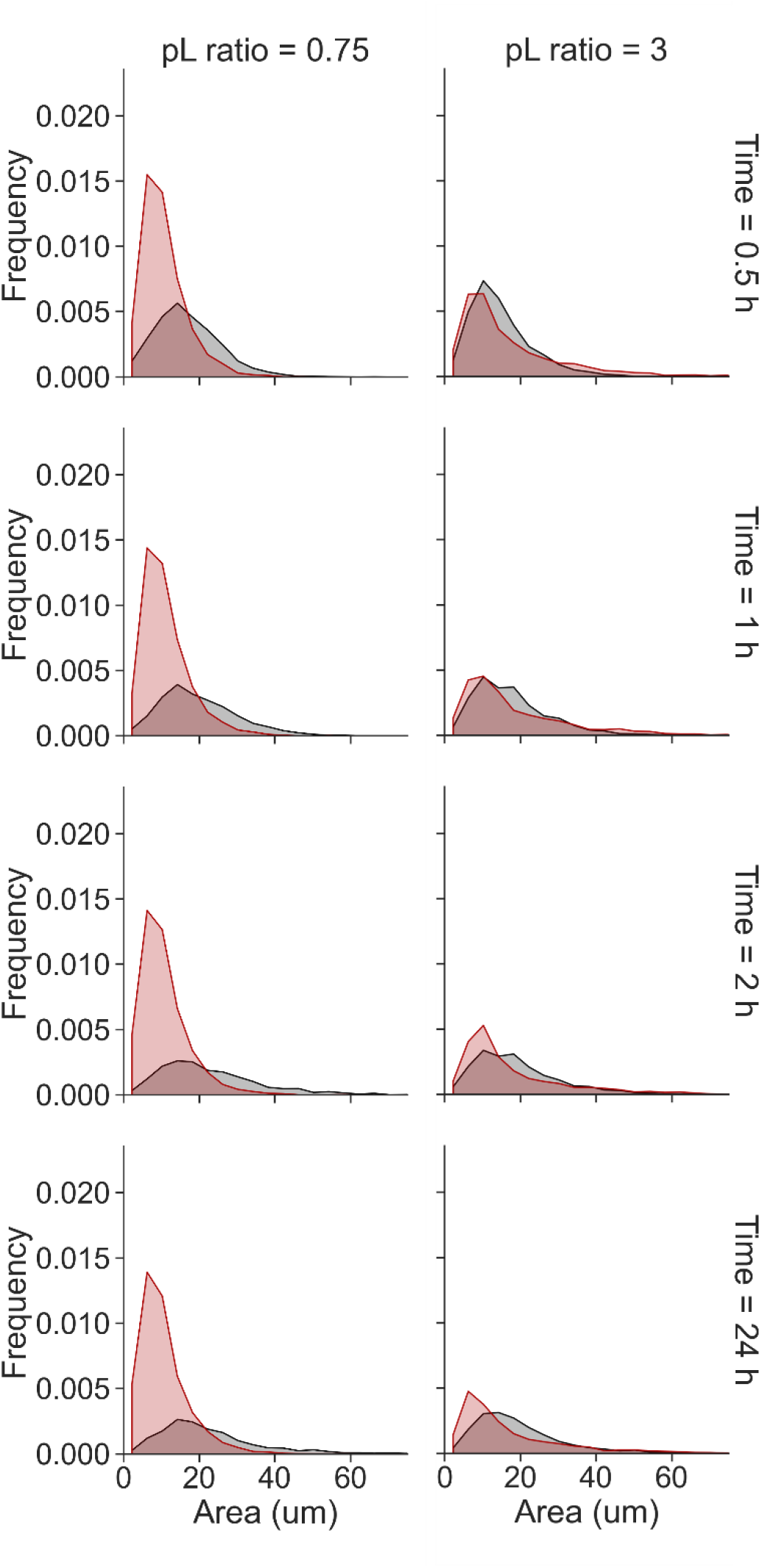
Frequency histograms area of Cy5-labelled coacervate droplet area over time. Droplets were from either 0.75:1 (Lys)_19-72_:RNA (left) or 3:1 (Lys)_5-24_:RNA (right) and contained active (red) or inactive (black) ribozyme.

**Figure 2 – supplement 4.**
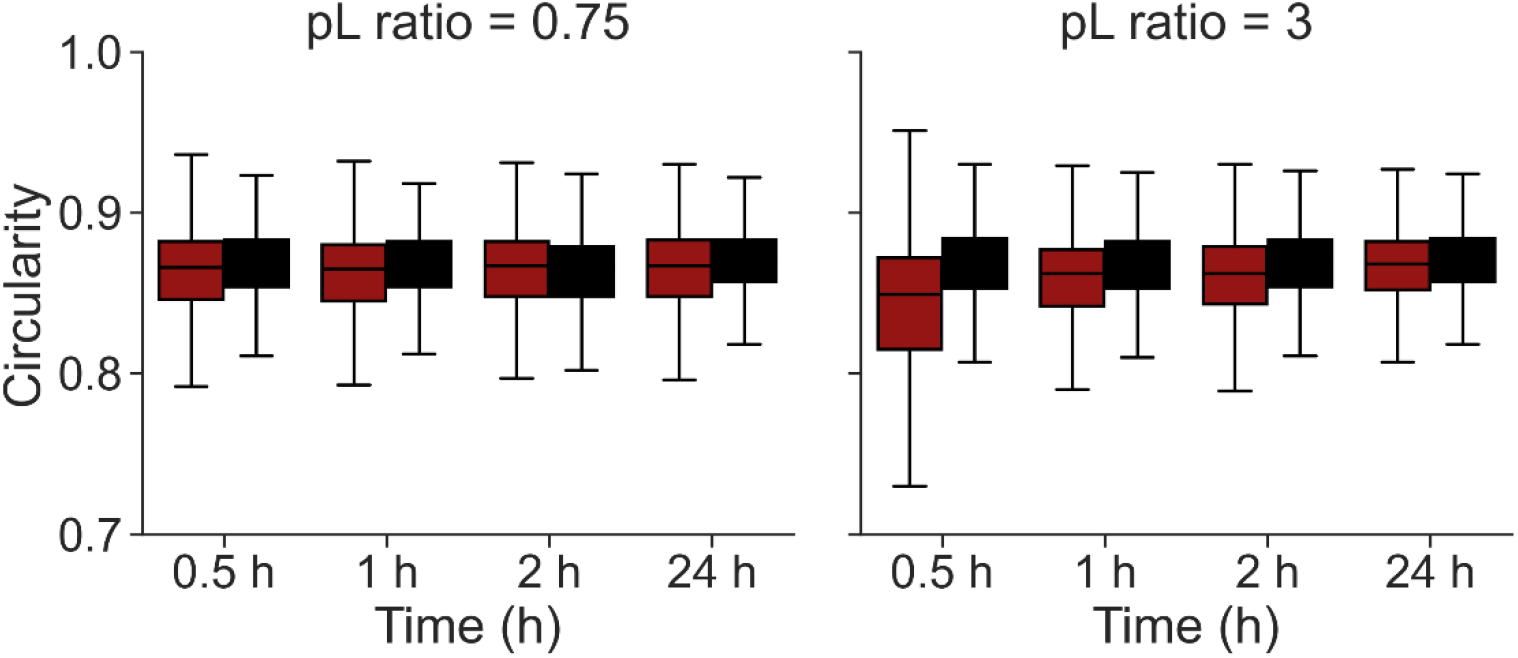
Circularity of Cy5-labelled coacervate droplets over time. Droplets were from either 0.75:1 (Lys)_19-72_:RNA (left) or 3:1 (Lys)_5-24_:RNA (right) and contained active (red) or inactive (black) ribozyme. Error bars are standard deviations.

**Figure 2 – supplement 5.**
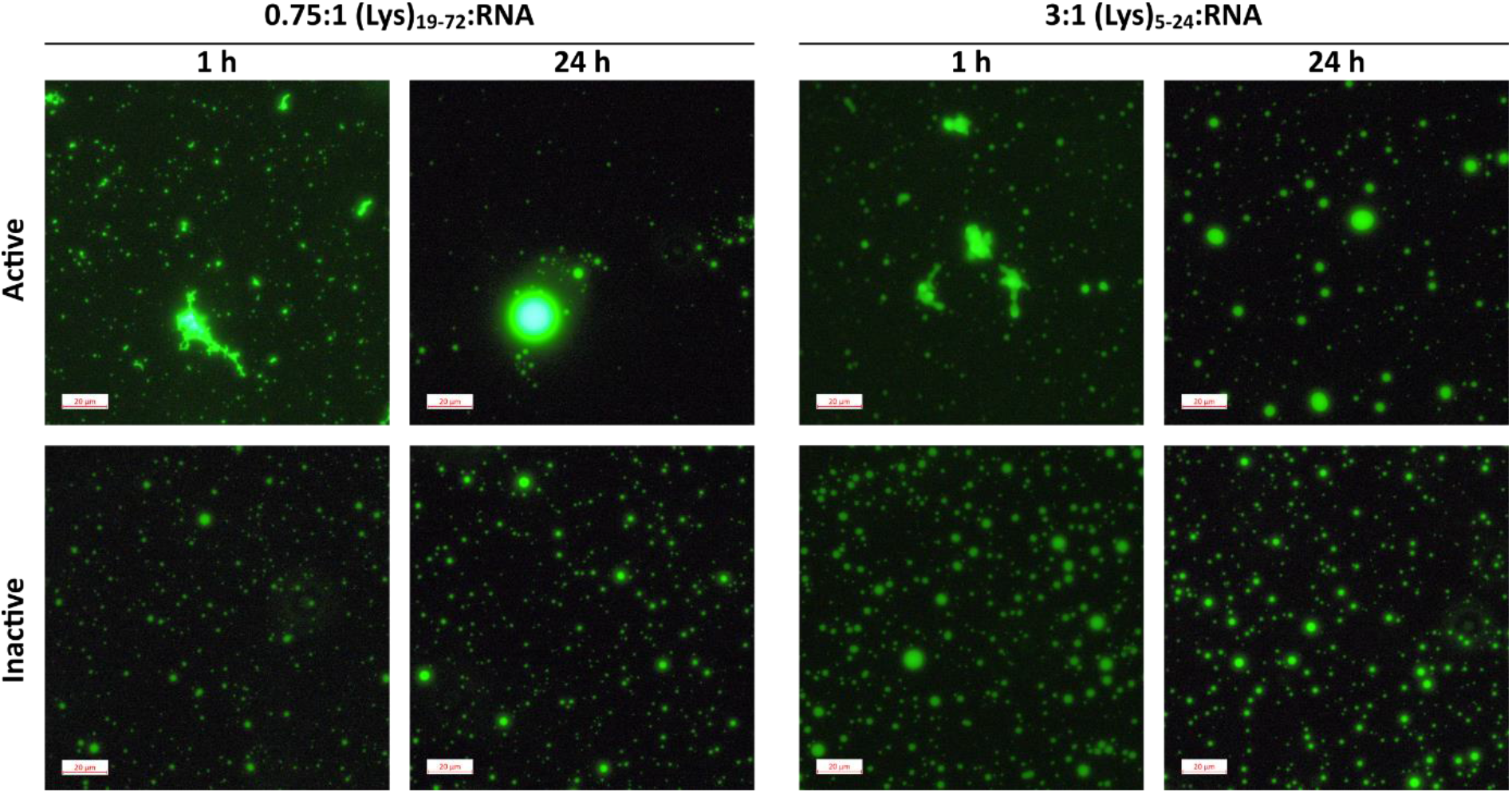
Comparison between condensates formed from pre-reacted E_L_ ribozyme and substrate and (Lys)_n_ at 1 h and 24 h after mixing. RNA and substrate were incubated for 2h at 30 °C with 10 mM MgCl_2_. (Lys)_n_ was added, then the samples were incubated for 24 h at 30 °C. Images were captured 1 h and 24 h after mixing.

**Figure 3 – supplement 1.**
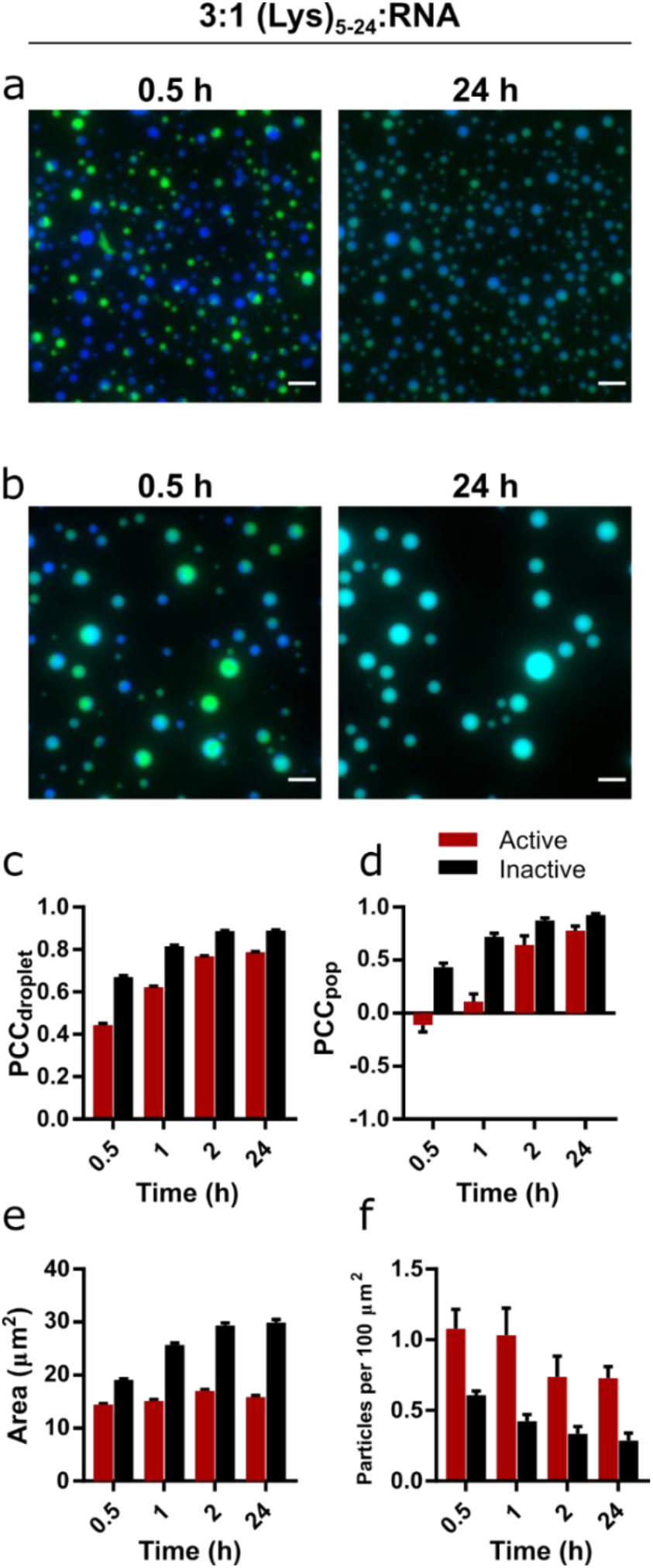
Mixing and content exchange between populations of active or inactive coacervate systems formed from R3C RNA and the (Lys)_5-24_ peptide. **a)** and **b)** Example images of mixtures of orthogonally labelled coacervate droplets prepared with 0.75:1 (Lys)_5-24_:RNA containing either active (**a**) or inactive (**b**). Each population in the set contained either 10 % FAM- or 10 % Cy5-tagged substrate (green and blue, respectively). The two populations in each set were mixed shortly after preparation and then imaged over 24 h in a passivated environment. All experiments were performed at 30 °C, pH 8.6 and 10 mM MgCl_2_ with a 1 mM total RNA monomer concentration and a 0.75:1 ratio of (Lys)_19-72_:RNA. The colocalization of the two fluorophores within single droplets is measured by the droplet Pearson coefficient (PCP_droplet_) **(c)**, whilst the colocalization of fluorophores in the overall population of droplets is measured by the population Pearson coefficient (PCC_pop_) **(d)**. The average particle area and number of particles per unit area over time are shown in **e** and **f** respectively. Particles were measured from at least 6 separate images. Error bars are standard errors. Scale bars = 10 μm.

**Figure 3 – supplement 2.**
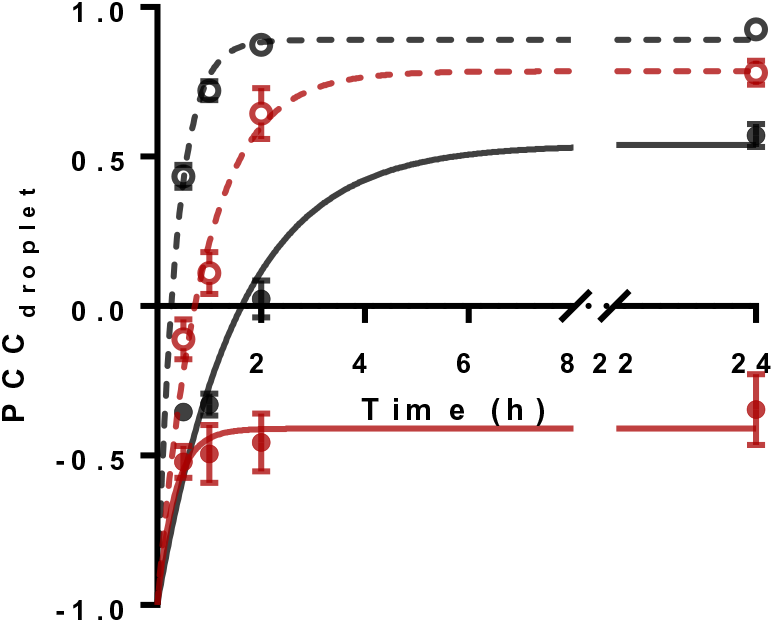
Comparison of mixing of labelled RNA in individual fusion droplets over time measured using the droplet Pearson correlation coefficient (PCC_droplet_). The plot displays a comparison between fluorophore colocalization over time in long (0.75:1 (Lys)_19-72_:RNA, solid lines) and short (3:1 (Lys)_5-24_:RNA) coacervate systems that contain either active (red) or inactive (black). All data are fitted with a single exponential. Error bars are standard errors.

**Figure 3 – supplement 3.**
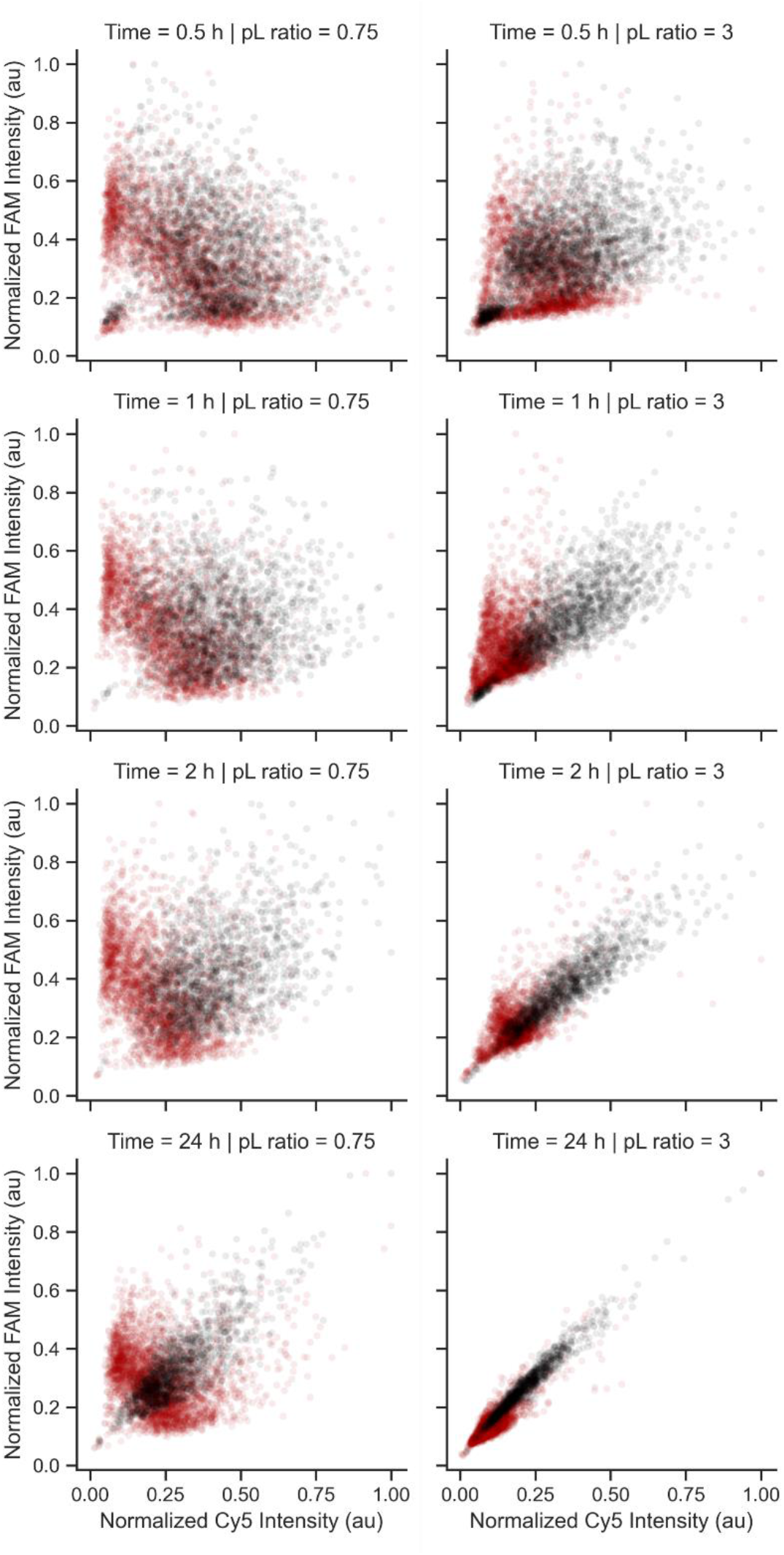
Intensity scatter plots showing the colocalization of FAM- and Cy5-labelled RNA in mixed droplet populations. Normalised Cy5 intensity is plotted on the X axis, whilst normalised FAM intensity is plotted on the Y axis. Each data point represents a single coacervate droplet. Droplets were from either 0.75:1 (Lys)_19-72_:RNA (left) or 3:1 (Lys)_5-24_:RNA (right) and contained active (red) or inactive (black) ribozyme. Data correspond to results reported in Figure 3 and S10.

## Source data

**Figure 1 – source data 1**. Unedited and uncropped gel image for **Figure 1b**, and labelled image showing key bands and conditions.

**Figure 1 – source data 2**. Numerical turbidity data for Figure 1c.

**Figure 1 – source data 3**. Unprocessed and uncropped fluorescence microscope image for 0.75:1 (Lys)_19-72_:RNA condensates (**Figure 1d**), imaged using 10 % Cy5-tagged substrate strand.

**Figure 1 – source data 4**. Unedited and uncropped gel images for ribozyme kinetics (**Figure 1e**). Lane identities are listed in the accompanying spreadsheet.

**Figure 1 – supplement 1 – source data 1**. Unedited and uncropped gel image for Figure 1 – supplement 1, and labelled image showing key bands and conditions.

**Figure 1 – supplement 2 – source data 1**. Unedited and uncropped gel image for Figure 1 – figure supplement 2, and labelled image showing key bands and conditions. Content identical to **Figure 1 – source data 2**.

**Figure 1 – supplement 2 – source data 2**. Numerical turbidity data. Content identical to **Figure 1 – source data 2**.

**Figure 1 – supplement 2 – source data 3**. Unprocessed and uncropped fluorescence microscope image for 3:1 (Lys)_5-24_:RNA condensates, imaged using 10 % Cy5-tagged substrate strand.

**Figure 1 – supplement 2 – source data 3**. Unedited and uncropped gel images for ribozyme kinetics (Figure 1e). Lane identities are listed in the included spreadsheet. Content identical to **Figure 1 – source data 2**.

**Figure 1 – supplement 3 – source data 1**. Unedited and uncropped gel image, and labelled image showing key bands and conditions.

**Figure 2 – source data 1**. Unprocessed and uncropped fluorescence microscope image for 0.75:1 (Lys)_19-72_:RNA condensates on passivated surfaces, imaged using 10 % Cy5-tagged substrate strand. Extracted numerical parameters are listed in the accompanying spreadsheet.

**Figure 2 – source data 2**. Unprocessed and uncropped fluorescence microscope image for 0.75:1 (Lys)_19-72_:RNA condensates on unpassivated surfaces, imaged using 10 % Cy5-tagged substrate strand. Extracted numerical parameters are listed in the accompanying spreadsheet.

**Figure 2 – supplement 1 – source data 1**. Unedited and uncropped gel image, and labelled image showing key bands and conditions.

**Figure 2 – supplement 2-4 – source data 1**. Unprocessed and uncropped fluorescence microscope image for 3:1 (Lys)_5-24_:RNA condensates on passivated and unpassivated surfaces, imaged using 10 % Cy5-tagged substrate strand. Extracted numerical parameters are listed in the accompanying spreadsheet.

**Figure 2 – supplement 5 – source data 1**. Unprocessed and uncropped fluorescence microscope image for 0.75:1 (Lys)_19-72_: prereacted RNA condensates, imaged using 10 % FAM-tagged substrate strand.

**Figure 2 – supplement 5 – source data 2**. Unprocessed and uncropped fluorescence microscope image for 3:1 (Lys)_5-24_: prereacted RNA condensates, imaged using 10 % FAM-tagged substrate strand.

**Figure 3 – source data 1**. Unprocessed and uncropped fluorescence microscope images of mixed populations of 0.75:1 (Lys)_19-72_:RNA condensates containing active ribozyme, imaged in both FAM and Cy5 channels. Extracted numerical parameters are listed in the accompanying spreadsheet.

**Figure 3 – source data 2**. Unprocessed and uncropped fluorescence microscope images of mixed populations of 0.75:1 (Lys)_19-72_:RNA condensates containing inactive ribozyme, imaged in both FAM and Cy5 channels. Extracted numerical parameters are listed in the accompanying spreadsheet.

**Figure 3 – supplement 1 – source data 1**. Unprocessed and uncropped fluorescence microscope images of mixed populations of 3:1 (Lys)_5-24_:RNA condensates containing active ribozyme, imaged in both FAM and Cy5 channels. Extracted numerical parameters are listed in the accompanying spreadsheet.

**Figure 3 – supplement 1 – source data 2**. Unprocessed and uncropped fluorescence microscope images of mixed populations of 3:1 (Lys)_5-24_:RNA condensates containing inactive ribozyme, imaged in both FAM and Cy5 channels. Extracted numerical parameters are listed in the accompanying spreadsheet.

**Figure 3 – supplement 2-3 – source data 1**. Numerical data for all plots displayed in Figure 3 – supplements 2 and 3.

## Notes

### Competing Interest Statement

The authors have declared no competing interest.

